# DNA-dependent macromolecular condensation drives self-assembly of the meiotic DNA break machinery

**DOI:** 10.1101/2020.02.21.960245

**Authors:** Corentin Claeys Bouuaert, Stephen Pu, Juncheng Wang, Dinshaw J. Patel, Scott Keeney

## Abstract

Formation of meiotic DNA double-strand breaks (DSBs) by Spo11 is tightly regulated and tied to chromosome structure, but the higher-order assemblies that execute and control DNA breakage are poorly understood. We address this question through molecular characterization of *Saccharomyces cerevisiae* RMM proteins (Rec114, Mei4 and Mer2)—essential, conserved components of the DSB machinery. Each subcomplex of Rec114–Mei4 (2:1 heterotrimer) or Mer2 (homotetrameric coiled coil) is monodisperse in solution, but they independently condense with DNA into dynamic, reversible nucleoprotein clusters that share properties with phase-separated systems. Multivalent interactions drive condensation, which correlates with DSB formation *in vivo*. Condensates fuse into mixed Rec114–Mei4–Mer2 clusters that further recruit Spo11 complexes. Our data show how the DSB machinery self-assembles on chromosome axes to create centers of DSB activity. We propose that multilayered control of Spo11 arises from recruitment of regulatory components and modulation of biophysical properties of the condensates.

## Introduction

During meiosis, Spo11-induced DNA double-strand breaks (DSBs) trigger a recombination program that promotes the pairing and subsequent segregation of homologous chromosomes (Hunter, 2015). Spo11 activity is highly controlled in terms of timing, number and distribution of DSBs, and is fundamentally integrated with the higher-order structure of meiotic chromosomes (Pan et al., 2011; Panizza et al., 2011; Lam and Keeney, 2015). However, the molecular mechanisms underlying the assembly of the DSB machinery and the multiscale levels over which Spo11 activity is controlled remain unclear (Keeney et al., 2014; Cooper et al., 2016).

In *S. cerevisiae*, DSB formation involves the coordinated action of ten proteins that can be classified into three subgroups (Lam and Keeney, 2015). A core complex of Spo11, Rec102, Rec104 and Ski8 forms the DSB enzyme that derives from an archaeal type II topoisomerase (Bergerat et al., 1997; Keeney et al., 1997; Robert et al., 2016a; Vrielynck et al., 2016) (Claeys Bouuaert *et al.*, in preparation). MRX (Mre11, Rad50 and Xrs2), a complex that is important for the processing of DSBs in mitotic and meiotic contexts, is also essential for DSB formation in yeast (Lam and Keeney, 2015).

Finally, the third component of the DSB machinery comprises the RMM proteins (Rec114, Mei4, Mer2). These proteins have been grouped together based on yeast-two-hybrid (Y2H) interactions, coimmunoprecipitation, and foci colocalization and interdependencies (Li et al., 2006; Maleki et al., 2007; Steiner et al., 2010; Miyoshi et al., 2012). However, in contrast to MRX and the Spo11 core complex, the formation of a stoichiometric RMM complex has not been demonstrated and the relationships between the proteins are unknown.

RMM lies at the crossroads between DSB formation and chromosome organization. The RMM proteins associate with chromatin early in meiotic prophase and form discrete, largely overlapping and interdependent foci along the chromosome axes (Henderson et al., 2006; Li et al., 2006; Maleki et al., 2007; Panizza et al., 2011). They physically interact with other components of the DSB machinery and the hotspot-targeting protein Spp1, thereby providing a connection between the chromosome axis and the sites of Spo11 cleavage (Arora et al., 2004; Maleki et al., 2007; Acquaviva et al., 2013; Sommermeyer et al., 2013). RMM proteins are conserved between yeast and other eukaryotes, including mice and humans, albeit with high sequence divergence (Kumar et al., 2015; Robert et al., 2016b; Stanzione et al., 2016; Tesse et al., 2017; Wang et al., 2019). However, their precise functions remain enigmatic. In particular, their biochemical properties and the mechanism whereby they participate in DSB formation are unknown.

To address this gap, we purified and characterized recombinant *S. cerevisiae* RMM proteins. We found that sub-complexes of Rec114– Mei4 and Mer2 independently form DNA-dependent macromolecular assemblies, which we refer to as condensates. The condensates can fuse and form structures that can further recruit the Spo11 core complex. By structure-function analysis, we show that protein-protein interactions within Rec114–Mei4, the DNA-binding activities of Rec114 and Mer2, and the interaction between Rec114 and the core complex subunits Rec102 and Rec104 are essential for RMM-promoted DSB formation. These findings reveal how the DSB machinery self-organizes into punctate DSB-competent clusters along meiotic chromosomes and highlight new potential mechanisms to control meiotic DSB formation.

## Results

### Rec114 and Mei4 assemble via their respective C and N-termini into a 2:1 complex

Despite the long known physical and functional relationships between Rec114, Mei4 and Mer2 (Arora et al., 2004; Li et al., 2006; Maleki et al., 2007), confirmed in other species including mice (Miyoshi et al., 2012; Stanzione et al., 2016; Tesse et al., 2017; Kumar et al., 2018), there is currently no evidence for the formation of a stoichiometric RMM complex. Indeed, foci colocalizations and functional dependencies between Rec114, Mei4 and Mer2 are incomplete, suggesting that they may not form an obligate three-subunit complex. To clarify these relationships and gain insights into their biochemical functions, we sought to purify a hypothetical tripartite Rec114–Mei4–Mer2 complex. However, while Mer2 alone and a two-subunit complex between Rec114 and Mei4 were readily purified (below), attempts to purify a three-subunit complex were unsuccessful (**Supplementary Figure 1A, B**). We propose that Rec114–Mei4 and Mer2 instead function as independent sub-complexes.

Rec114 is 428 amino acids, much of it predicted to be disordered (residues ∼150 to 400, **Figure 1A**, top). The N-terminal 140 amino acids contain six conserved signature sequence motifs (SSMs) and a seventh is located at its C terminus (Maleki et al., 2007; Kumar et al., 2010; Tesse et al., 2017). Recent structural analyses of the N-terminal domain of *Mus musculus* REC114 showed that the SSMs are part of secondary structures that form the core of a pleckstrin homology (PH)-like fold (Kumar et al., 2018; Boekhout et al., 2019). In contrast to Rec114, Mei4 is predicted to be mostly ordered (**Figure 1A**, bottom). Six SSMs were identified in Mei4, including two within the N-terminal 50 amino acids (Kumar et al., 2010).

**Figure 1:**
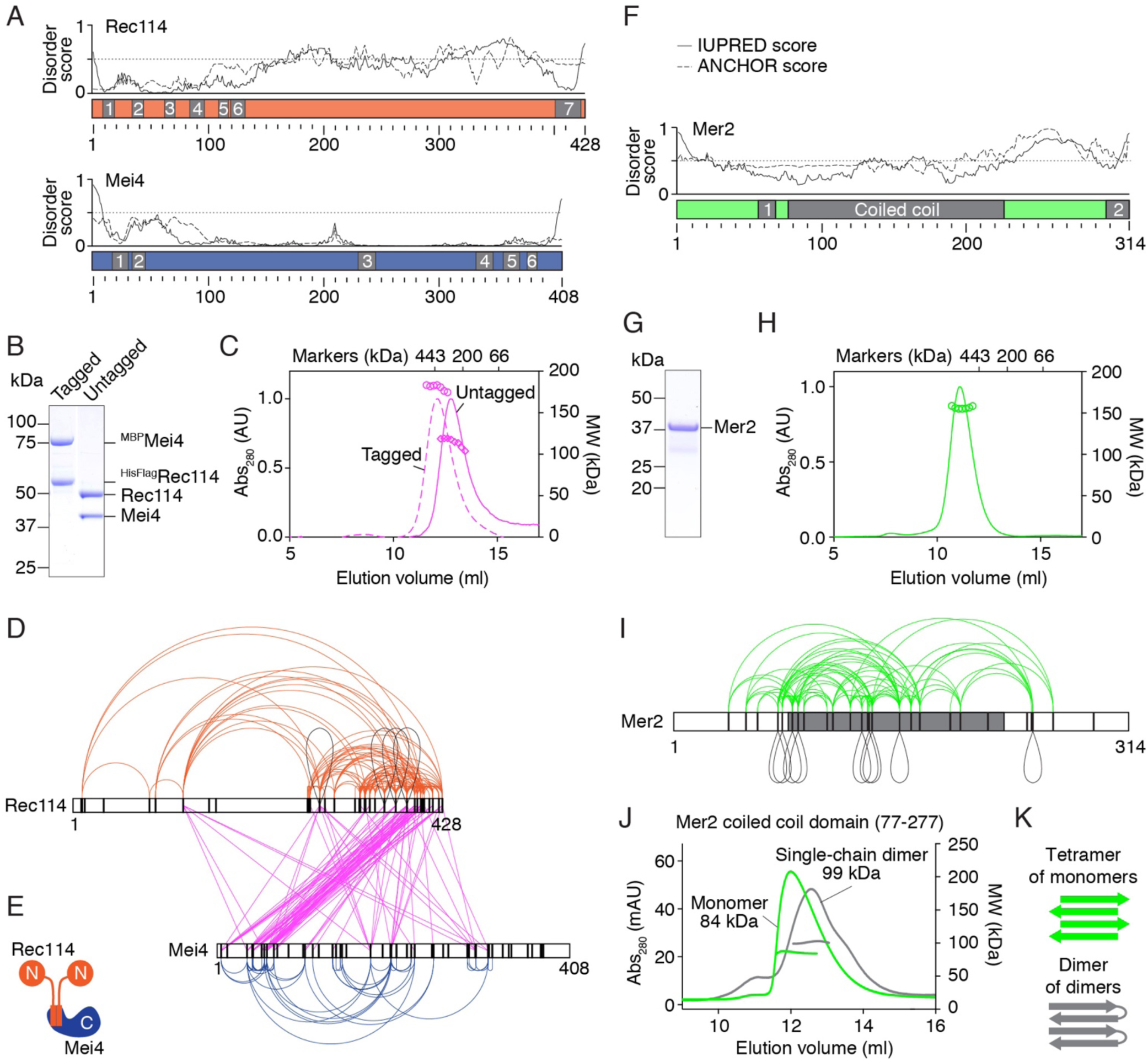
Purification and subunit arrangement of the *S. cerevisiae* RMM proteins. A, B. Prediction of protein disorder using the IUPRED server (Dosztanyi, 2018). The ANCHOR score predicts the transition from unstructured to structured depending on a binding partner. Previously identified SSMs are highlighted (Kumar et al., 2010; Tesse et al., 2017). B. SDS-PAGE of purified tagged and untagged Rec114–Mei4 complexes. 4 µg was loaded. C. SEC-MALS analysis of tagged and untagged Rec114–Mei4. The traces show UV absorbance (left axis), circles are molar mass measurements across the peak (right axis). Elution positions of protein standards are marked. D. XL-MS analysis of Rec114–Mei4 (4812 crosslinked peptides, 258 distinct crosslinked pairs of lysines). Black loops are intermolecular self-links. Black vertical lines indicate lysines. E. Cartoon of the Rec114–Mei4 complex. See also **Supplementary Figure 1**. F. Protein disorder prediction for Mer2. The predicted coiled coil and previously identified SSMs are highlighted (Kumar et al., 2010; Tesse et al., 2017). G. SDS-PAGE of purified Mer2. 4 µg was loaded. H. SEC-MALS analysis Mer2. I. XL-MS analysis of Mer2 (487 crosslinked peptides, 89 distinct crosslinked pairs of lysines). J. SEC-MALS analysis of the coiled coil domain of Mer2 and a single-chain dimer variant of the coiled coil domain. A tetramer of monomers and a dimer of single-chain dimers both have an expected MW of 70 kDa. The difference between the profiles of the monomer and single chain dimer can be explained by reduced degrees of freedom (tension) in the single-chain dimer and heterogeneity. K. Interpretive cartoon of the molecular arrangement of the coiled coil domain of Mer2.

We purified tagged and untagged versions of Rec114–Mei4 from baculovirus-infected insect cells and analyzed the complexes by size-exclusion chromatography followed by multi-angle light scattering (SEC-MALS, **Figure 1B, C**). This revealed molar masses (MW) of 180 kDa and 114 kDa for the tagged and untagged complexes, respectively. These results and the intensities of Coomassie-stained bands suggest that the complex might contain two Rec114 subunits and one Mei4 subunit (expected sizes 200 kDa and 146 kDa for tagged and untagged, respectively). This is supported by a 2:1 ratio of mass spectrometry spectral counts (**Supplementary Figure 1C**).

To delineate the molecular arrangement of Rec114 and Mei4, we analyzed purified complexes by crosslinking coupled to mass spectrometry (XL-MS). This analysis yielded 258 distinct pairs of crosslinked lysine residues (**Figure 1D & Supplementary Table 1**). The C-terminus of Rec114 crosslinked extensively to the N-terminus of Mei4 (pink lines), implying that these are the primary interaction regions. In addition, four inter-molecular self-links (positions where two identical lysines are crosslinked) were found toward the C-terminal domain of Rec114 (black loops in **Figure 1D**). This result supports the 2:1 stoichiometry and shows that this domain likely homo-dimerizes (**Figure 1E**).

Truncated polypeptides that retained SSM7 of Rec114 and SSMs 1 and 2 of Mei4 (fragments Rec114^(375-428)^ and Mei4^(1-43)^) formed a complex with a 2:1 stoichiometry (**Supplementary Figure 1D-H**). C-terminal fragments of Rec114 alone formed dimers, thus Rec114 dimerization does not depend on Mei4 (**Supplementary Figure 1H**). Mutation of a conserved residue of Rec114 (F411A) abolished dimerization, which disrupted the interaction with Mei4 similarly to an equivalent mutation in the *S. pombe* Rec114 ortholog, Rec7 (**Supplementary Figure 1I-K**) (Steiner et al., 2010). The Rec114-F411A mutation does not compromise protein expression *in vivo*, but it abolishes the formation of chromatin-associated Rec114 foci and Spo11-mediated DSBs, leading to spore death (**Supplementary Figure 1L-O**).

### Mer2 forms a homotetrameric parallel-antiparallel α-helical bundle

Mer2 is a 314 amino-acid protein with a predicted coiled coil (spanning residues ∼77-227) and two conserved SSMs, including one at its C-terminus (Engebrecht et al., 1991; Tesse et al., 2017). The region connecting the coiled coil and SSM2 is predicted to be disordered (**Figure 1F**). SEC-MALS of untagged Mer2 purified from *E. coli* revealed a 156 kDa species, consistent with a tetramer (143 kDa expected) (**Figure 1G, H**). However, the elution volume on the size-exclusion column was consistent with a considerably larger complex, suggesting a highly elongated shape (compare with the position of the markers in **Figure 1H**).

XL-MS revealed nine intermolecular self-links, providing further evidence for multimerization (**Figure 1I**). The self-links occurred all along the predicted coiled-coil domain, consistent with a parallel arrangement of α-helices, but long-range crosslinks were also detected along the coiled coil. If the coiled coil forms uninterrupted helices, crosslinks between residues further than about 18 amino acids cannot be explained by intra-molecular events or by intermolecular events between parallel coiled coils. Therefore, a likely possibility is that the coiled coil contains both parallel and antiparallel helices.

To address this, we first analyzed the coiled-coil domain of Mer2 alone (residues 77-227) by SEC-MALS and found that it is tetrameric, thus the N and C-termini are dispensable for tetramerization (**Figure 1J**). Next, we engineered a single-chain dimer in which two copies of the coiled-coil domain are separated by a 19 amino-acid linker, too short to allow a parallel intramolecular coiled coil. SEC-MALS analysis showed that this construct assembles a complex of similar size as the monomeric construct (99 vs. 84 kDa), consistent with two single-chain dimers, each one folded in antiparallel (**Figure 1J, K**). Alternative scenarios all predict that the single-chain dimer would be artificially elongated, which would lead to a faster elution from the size exclusion column. However, the construct eluted at a similar volume as the monomer, if later, suggesting it is somewhat constrained. The single-chain dimer peak was slightly broader, probably due to some heterogeneity. Mer2 therefore likely assembles as a homotetrameric alpha-helical bundle with two pairs of parallel helices arranged in an anti-parallel configuration (**Figure 1K**).

### Rec114–Mei4 and Mer2 form DNA-induced condensates

*S. cerevisiae* Rec114, Mei4 and Mer2 form chromatin-associated foci, similar to their homologs in *S. pombe* (Rec7, Rec24, Rec15) and mice (REC114, MEI4 and IHO1) (Henderson et al., 2006; Li et al., 2006; Lorenz et al., 2006; Maleki et al., 2007; Kumar et al., 2010; Bonfils et al., 2011; Panizza et al., 2011; Kumar et al., 2018). However, the physical nature of these foci is unclear, and so is the relationship between these structures, the biochemical properties of Rec114, Mei4 and Mer2, and their DSB-promoting functions.

First, we set out to address whether Rec114–Mei4 and Mer2 bind DNA. We titrated the complexes in the presence of 20-, 40-, and 80-bp radiolabeled substrates and analyzed DNA binding by native gel electrophoresis (**Figure 2A, B**). Rec114–Mei4 and Mer2 bound all three DNA substrates, and the binding affinity increased with substrate size (**Figure 2A, B, C, E**). This preference for longer substrates was confirmed in a competition assay using a labeled 80-bp substrate in the presence of unlabeled 20-bp or 80-bp competitors. The 20-bp substrate provided a poor competitor, in contrast to the 80-bp substrate (**Figure 2D, F, Supplementary Figure 2A, B**). Titrations of Rec114–Mei4 and Mer2 revealed well-shifts with no discrete migrating bands, and remarkable switch-like transitions from no binding to complete binding within narrow (2 to 4-fold) ranges, suggesting the cooperative assembly of higher-order structures (**Figure 2A, B, C, E**).

**Figure 2:**
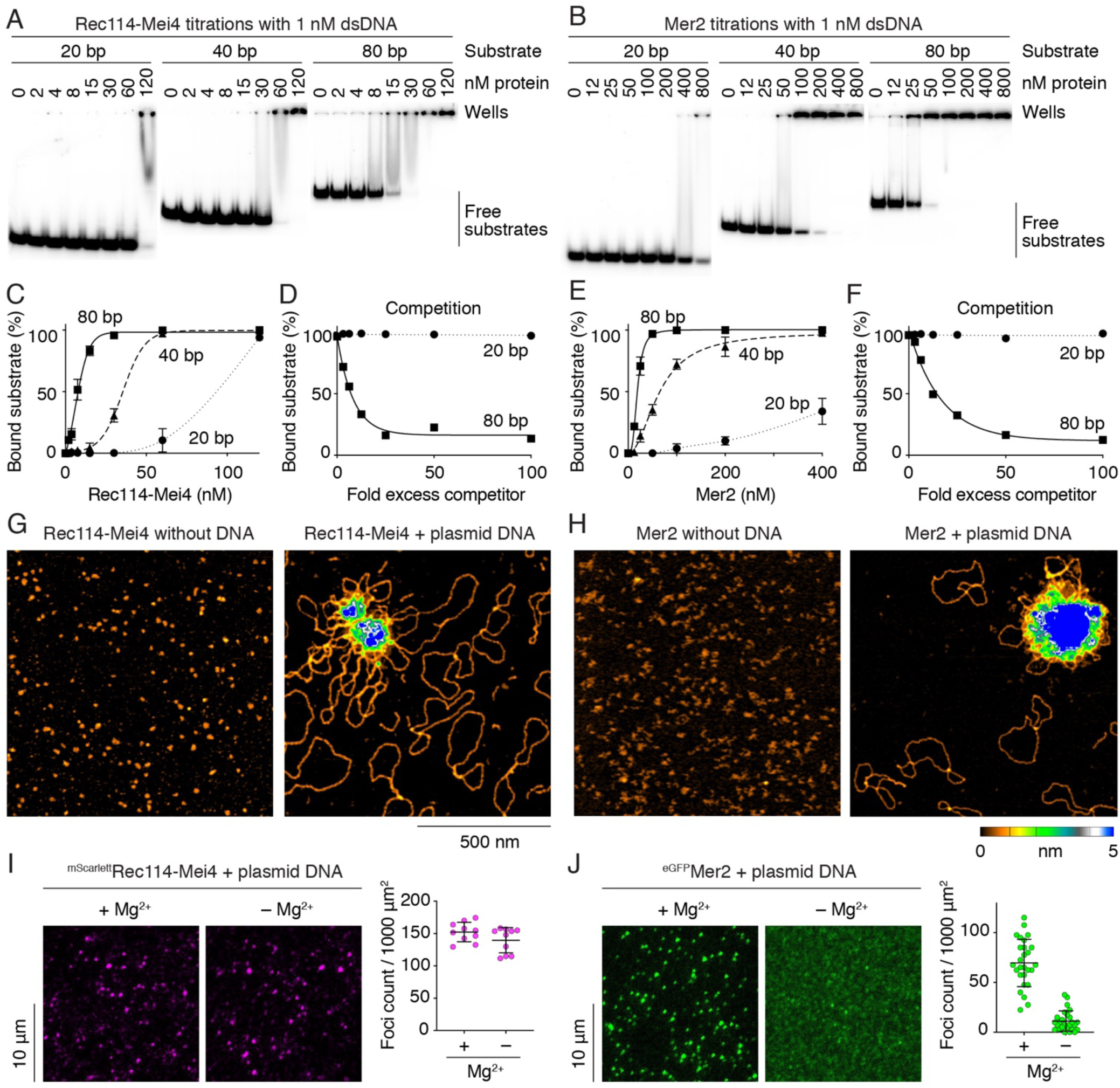
Rec114–Mei4 and Mer2 form condensates on DNA. A, B. Gel shift analysis of Rec114–Mei4 (A) or Mer2 (B) binding to 20-, 40- or 80 bp DNA substrates. C, E. Quantification of the gel-shift analyses in panels A and B, respectively. Error bars are ranges from two independent experiments. Lines are sigmoidal curves fit to the data, except for the 20 bp substrate (smooth spline fits). The apparent affinities of Rec114–Mei4 for the DNA substrates are: 6 ± 1.4 nM (80 bp, mean and range); 35 ± 1.3 nM (40 bp); ≈ 80 nM (20 bp). The apparent affinities of Mer2 for the DNA substrates are: 19 ± 1.5 nM (80 bp); 64 ± 15 nM (40 bp); > 400 nM (20 bp). Here and elsewhere, concentrations for Rec114–Mei4 refer to the trimeric complex, but for Mer2 they refer to the monomer. Therefore, even though the complexes appear to have very different affinities for DNA, they are in fact comparable if the basal units (i.e. trimers and tetramers, respectively) are considered. D, F. Competition assay of Rec114–Mei4 (D) or Mer2 (F) binding to an 80 bp radiolabeled DNA substrate (1 nM) in the presence of 20 bp or 80 bp cold competitor substrates. Fold excess is normalized to amounts in nucleotides. Lines are one-phase decay fits to the data. G, H. AFM imaging of 12 nM Rec114–Mei4 (G) and 50 nM Mer2 (H) in the absence (left) or in the presence (right) of 1 nM plasmid DNA (pUC19). I, J. Visualization of nucleoprotein condensates by epifluorescence microscopy using tagged Rec114–Mei4 (I) or Mer2 (J) in the presence or absence of MgCl_2_. Foci were defined using a fixed intensity threshold between samples. Each point represents the measurement from a field of view. Error bars show mean ± SD from 8–10 fields of view.

To directly visualize the DNA-bound particles, we turned to atomic force microscopy (AFM). In the absence of DNA, Rec114–Mei4 and Mer2 formed small, relatively homogeneous particles on the mica surface (**Figure 2G, H**, left panels). In contrast, the presence of plasmid DNA caused Rec114–Mei4 and Mer2 to assemble into large nucleoprotein structures with DNA loops emanating from protein-dense clusters (**Figure 2G, H**, right panels). The majority of the plasmid molecules remained unbound and the background was devoid of particles from individual free protein complexes, showing that this clustering property is extremely cooperative. The size of the clusters (∼0.2 µm diameter for Rec114–Mei4 and ∼0.4 µm for Mer2) indicates that these must contain many hundreds of proteins.

Given the clusters’ size, we reasoned that we should be able to directly visualize them by fluorescence microscopy. As expected, a Rec114–Mei4 complex that carried an mScarlet fluorophore fused to the N-terminus of Rec114 yielded bright foci in the presence of DNA (**Figure 2I**). These foci were not dependent on the presence of magnesium ions. An eGFP-tagged Mer2 complex also produced DNA-dependent foci in the presence of Mg^2+^, but gave only diffuse fluorescence signal without Mg^2+^ (**Figure 2J & Supplementary Figure 2C, D**).

### Rec114–Mei4 and Mer2 nucleoprotein condensates share properties with systems that undergo phase transition

The Rec114–Mei4 and Mer2 nucleoprotein clusters are reminiscent of biomolecular condensates that form intracellular membrane-less compartments and control a variety of biological processes, including transcription, signal transduction and stress responses (Li et al., 2012; Su et al., 2016; Wheeler et al., 2016; Banani et al., 2017; Boulay et al., 2017; Boeynaems et al., 2018). Phase separation of such biomolecules is often driven by combinations of weak interactions between multivalent components (Li et al., 2012; Banani et al., 2017; Boeynaems et al., 2018). These systems typically share certain biophysical properties: condensates tend to be reversible, are promoted by molecular crowding, can fuse with one another, and may undergo sol-gel transitions over time as the condensates accumulate entanglements or stabilize in a low-entropy form.

We therefore characterized DNA-dependent Rec114–Mei4 and Mer2 condensates to address whether they display behaviors typical of phase-separated systems. To do this, we generated functional Rec114– Mei4 and Mer2 variants covalently coupled to Alexa594 and Alexa488 fluorophores, respectively, to minimize potential steric effects or oligomerization of bulky fluorescent protein tags (**Supplementary Figure 2E-H**).

We first consider the effect of molecular crowding. Indeed, addition of the crowding agent polyethylene glycol (5% PEG-8000) dramatically increased the intensity of the condensates for both Rec114–Mei4 and Mer2 (**Figure 3A, B**). Protein titrations revealed complex, sometimes counter-intuitive behaviors, including a decrease in focus numbers with increasing protein concentrations (**Supplementary Figure 3A, B**). These behaviors likely reflect balances between nucleation, growth, and collapse of the condensates (see legend to **Supplementary Figure 3A,B**).

**Figure 3:**
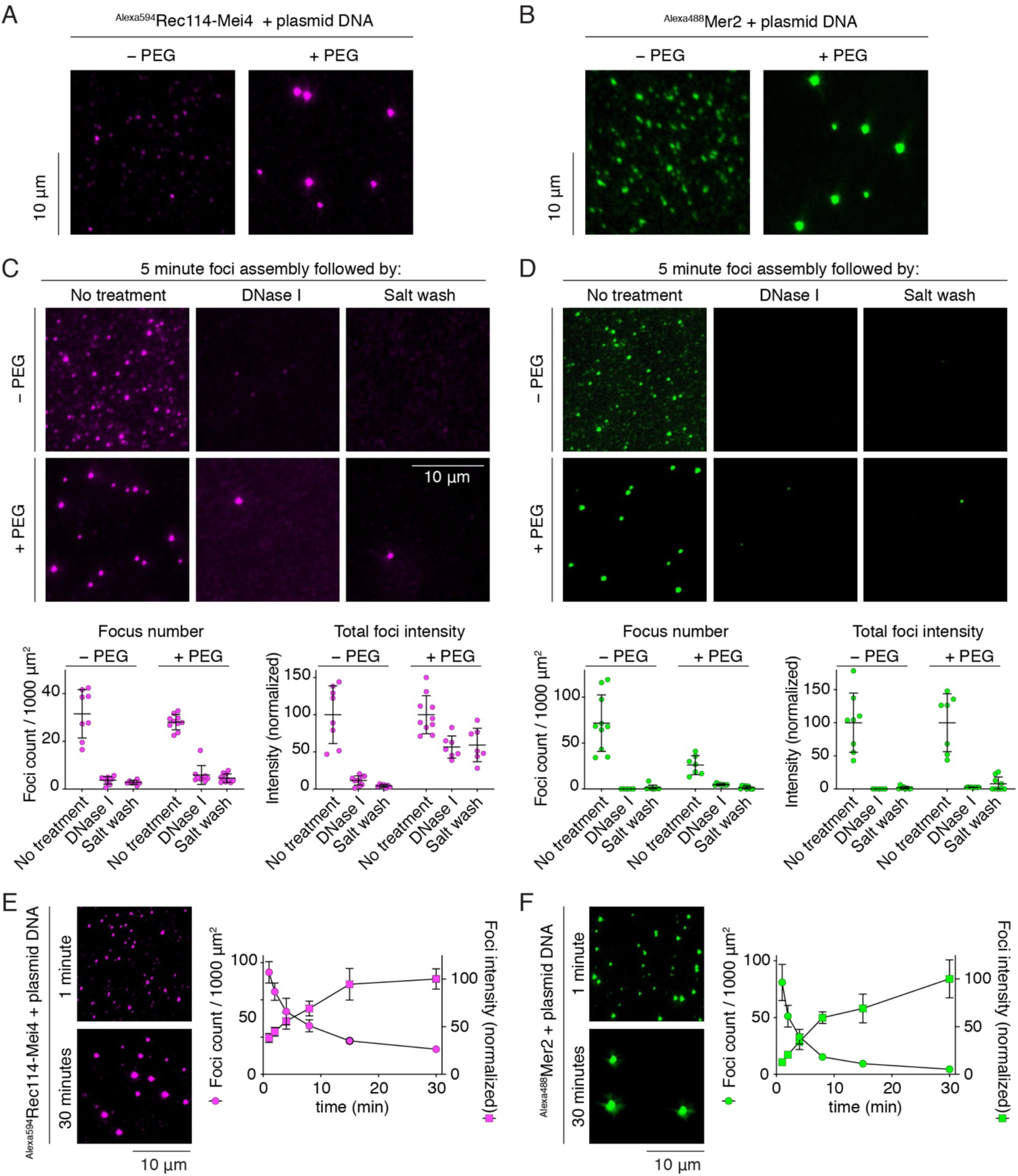
Properties of Rec114–Mei4 and Mer2 nucleoprotein condensates. A, B. Effect of a crowding agent (PEG) on formation of nucleoprotein condensates visualized using covalently fluorophore-labeled Rec114–Mei4 (A) or Mer2 (B). C, D. Effect of challenging Rec114–Mei4 (C) or Mer2 (D) nucleoprotein condensates with DNase I or 0.5 M NaCl. Condensates were assembled for 5 minutes prior to challenge. Quantification is provided of focus numbers per 1000 µm^2^ and of the total fluorescence intensity within foci within fields of view (normalized to mean of the no-treatment controls). Error bars show mean ± SD from 8–10 fields of view. E, F. Time course of the assembly of foci of Rec114–Mei4 (A) or Mer2 (B) in the presence of plasmid DNA. The x axis indicates the time in solution before plating, upon which DNA is immobilized to the glass slide while soluble protein is still free to diffuse. Quantification is provided of focus numbers and average focus intensity (normalized to the mean at 30 min). Error bars show mean ± SD from 8–10 fields of view.

Condensation was inhibited at high ionic strength, suggesting that condensates depend on electrostatic interactions, probably between the negatively charged DNA backbone and positively charged protein residues (**Supplementary Figure 3C, E**). In addition, competition experiments revealed preferential incorporation of larger DNA molecules, consistent with the hypothesis that condensation is driven by multivalency of the substrate (**Supplementary Figure 3G-J**).

To address whether the nucleoprotein condensates are reversible, we assembled Rec114–Mei4 and Mer2 condensates for 5 minutes in the presence or absence of PEG and then challenged them by treatment with DNase I or 500 mM NaCl. In the absence of PEG, DNase I and salt treatments almost completely dissolved the condensates, showing that they are indeed reversible (**Figure 3C, D**). However, in the presence of PEG, about half of the condensate-associated fluorescence signal of Rec114–Mei4 resisted DNase I and salt treatments. This result suggested that Rec114–Mei4 condensates can adopt to a more stable form. We found that the reversibility of Rec114–Mei4 condensates decreased over time, accentuated by molecular crowding (**Supplementary Figure 3D**). With a short assembly time, Mer2 condensates were unable to resist dissolution whether PEG was present or not (**Figure 3D**), but longer incubation times with PEG allowed Mer2 as well to form relatively resistant foci (**Supplementary Figure 3F**). These results suggest that condensates of Rec114–Mei4 and, to a lesser extent, Mer2 may spontaneously mature into irreversible, perhaps gel-like, structures, as has been observed for other phase separation systems (Lin et al., 2015; Patel et al., 2015; Xiang et al., 2015; Banani et al., 2017).

Finally, we sought to better understand how the condensates assemble, envisioning several scenarios: (i) Nucleation could be limiting, with focus growth resulting principally from incorporation of protein from soluble pools. (ii) Frequent nucleation events could occur at early time points leading to large numbers of small foci, whereupon some foci dissolve and others grow. (iii) Finally, frequent nucleation events could occur leading to numerous small foci that then collide and fuse to yield fewer, larger foci (**Supplementary Figure 4A**). To distinguish between these possibilities, we devised an experiment where DNA was immobilized at varied time points by spreading the assembly reactions on glass slides. This should prevent foci fusion, but not subunit exchange with soluble protein pools. Images were then captured at a late time point (>1 hour), so the time variable in this experiment is the period that the DNA is free in solution before constraint. In the first two assembly models, foci grow by addition from soluble protein pools, so they predict that DNA immobilization should have no effect and that all the reactions should be identical. In contrast, if fusion is a major growth mechanism, the experiment would reveal decreasing numbers and increasing intensities of foci over time. The latter outcome is what we observed for both Rec114–Mei4 and Mer2 (**Figure 3E, F**), thus focus growth in this assay is determined largely by fusion. However, the rates of dynamic exchanges from soluble pools are likely to be a function of experimental conditions, and such exchange may be particularly important *in vivo* where viscous drag on chromosomes may inhibit fusion.

### DNA binding is important for Rec114–Mei4 and Mer2 function

By truncation analysis, we found that the C-terminal domain of Rec114 is necessary and sufficient for DNA binding (**Supplementary Figure 4B**). Alanine substitution of four positively charged residues within this domain yielded a Rec114–Mei4 complex (referred to as 4KR) with reduced DNA-binding activity (**Figure 4A**). Similarly, alanine substitutions within a conserved patch of basic residues located towards the C-terminus of Mer2 (KRRR) yielded a DNA-binding defective mutant (**Figure 4B, Supplementary Figure 4C**). As expected if multivalent protein-DNA interactions contribute to condensate formation, both the Rec114-4KR and the Mer2-KRRR mutant proteins showed strongly reduced focus formation *in vitro* (**Figure 4C, D**).

**Figure 4:**
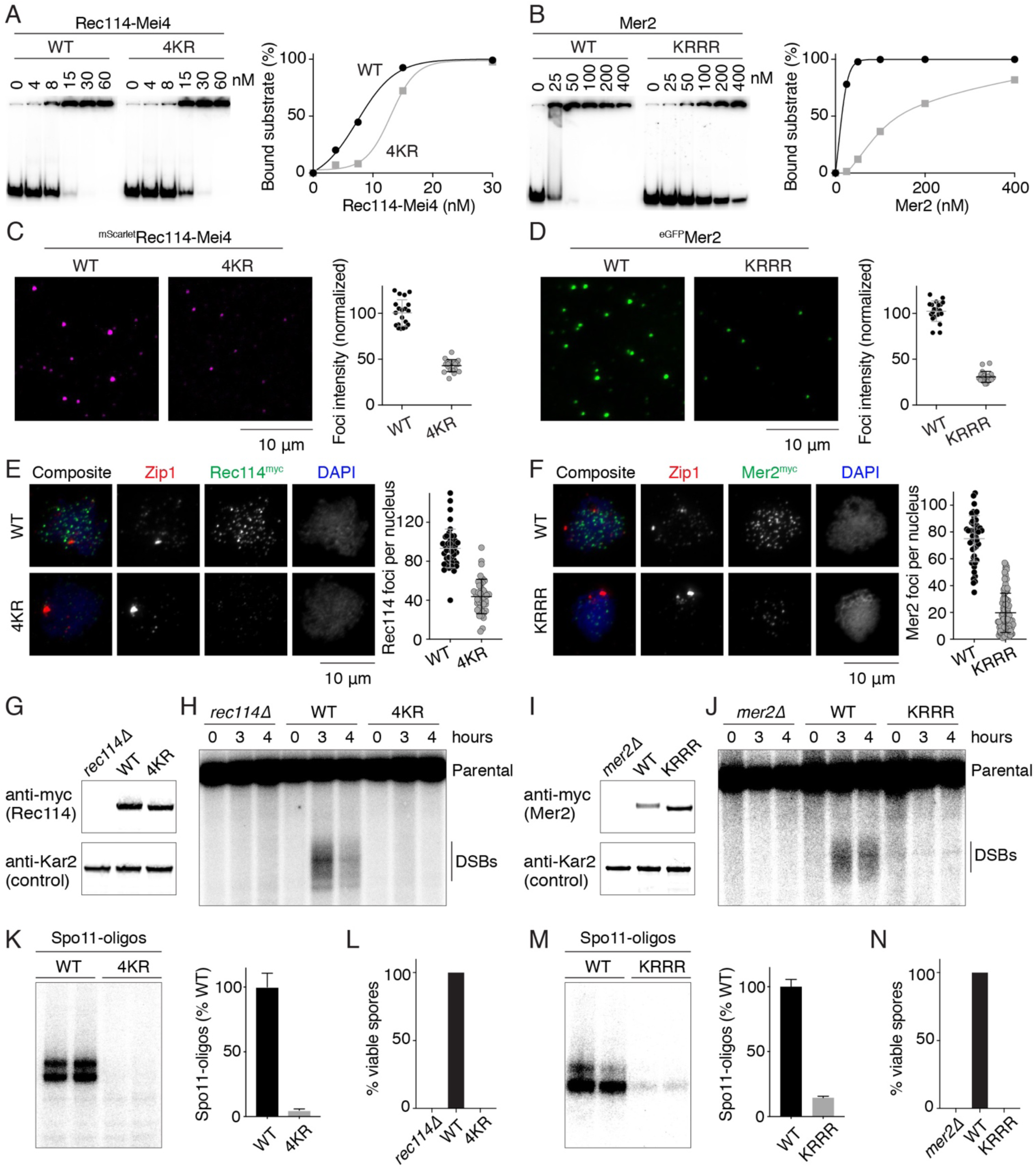
DNA binding by Rec114–Mei4 and Mer2 is important for their macromolecular condensation *in vitro* and *in vivo* and for Spo11-induced break formation. A, B. DNA-binding of wild-type (WT) and mutant Rec114–Mei4 (A) or Mer2 (B) complexes to an 80-bp DNA substrate measured by gel shift assay. The Rec114-4KR mutant has four positively charged residues within the C terminus of Rec114 (R395, K396, K399, R400) mutated to alanine. The Mer2-KRRR mutant has four positively charged residues located towards the C terminus of Mer2 (K265, R266, R267, R268) mutated to alanine. Lines on graphs are sigmoidal curves fit to the data, except for the Mer2-KRRR mutant data that is a smooth spline fit. C, D. Effect of the Rec114-4KR (C) or Mer2-KRRR mutations (D) on condensation *in vitro*. Reactions included 5% PEG. Each point is the average of the intensities of foci in a field of view, normalized to the overall mean for wild type. Error bars show mean ± SD. E, F. Immunofluorescence on meiotic chromosome spreads for myc-tagged Rec114 (E) or Mer2 (F). The number of foci per leptotene or early zygotene cell is plotted. Error bars show mean ± SD. G, I. Western blotting of meiotic protein extracts for Rec114 (G) or Mer2 (I). H, J. Southern blot analysis of meiotic DSB formation at the *CCT6* hotspot in wild type and mutant Rec114 (H) and Mer2 (J) strains. K, M. Labeling of Spo11-oligo complexes in wild type and mutant Rec114 (K) and Mer2 (M) strains. Error bars represent the range from two biological replicates. L, N. Spore viability of wild type and mutant Rec114 (L) and Mer2 (N) strains.

To establish the *in vivo* consequences of attenuating these DNA-binding activities, we mutated the DNA-binding residues (4KR and KRRR) in the context of yeast strains that carried myc-tagged Rec114 and Mer2, respectively. Wild-type Rec114^myc^ and Mer2^myc^ formed foci on meiotic chromosomes, as revealed by immunofluorescence staining of meiotic nuclear spreads (**Figure 4E, F**) (Henderson et al., 2006; Li et al., 2006; Maleki et al., 2007). In contrast, the *rec114-4KR* and the *mer2-KRRR* mutants formed much fewer foci. This could not be attributed to destabilization of the proteins because immunoblot analyses showed that both mutants were expressed at similar levels as wild type (**Figure 4G, I**). In the case of Mer2, the KRRR mutation caused the protein to accumulate and persist longer during meiosis (**Supplementary Figure 4D**). The Mer2-KRRR protein also had a higher electrophoretic mobility than the wild-type, probably because it failed to become phosphorylated. It therefore appears that DNA-binding by Mer2 is a prerequisite for Mer2 phosphorylation, which is known to promote turnover of the protein (Henderson et al., 2006).

If DNA-driven condensation by Rec114–Mei4 and Mer2 is indeed important for their biological function, the Rec114-4KR and Mer2-KRRR mutations would be expected to confer defects in meiotic DSB formation. This was indeed the case: Southern blot analysis at the DSB hotspot *CCT6* showed that both mutants were defective for DSB formation (**Figure 4H, J**). Furthermore, levels of Spo11-oligo complexes were also reduced, indicating that this was not a locus-specific effect (**Figure 4K, M**). As anticipated, these DSB defects caused low spore viability (**Figure 4L, N**). In conclusion, the DNA-binding activities of Rec114–Mei4 and Mer2 are essential for DNA-driven condensation *in vitro* and *in vivo* and for their DSB-promoting activity.

### Rec114–Mei4 and Mer2 form joint nucleoprotein condensates

*In vivo*, Rec114, Mei4 and Mer2 form partially overlapping foci (Li et al., 2006; Maleki et al., 2007) and yield coincident ChIP signals (Panizza et al., 2011). We therefore hypothesized that they may function together as joint condensates. We tested this idea by mixing fluorescent Rec114–Mei4 and Mer2 either before or after DNA-driven condensation (**Figure 5**). Premixing Rec114–Mei4 and Mer2 prior to DNA-driven condensation led to mixed foci with essentially perfect overlap between the two components (**Figure 5A**). Colocalization was evident even with a large excess of DNA, thus formation of joint foci was not driven by independent assemblies on a limiting number of substrate molecules (**Supplementary Figure 6**).

**Figure 5:**
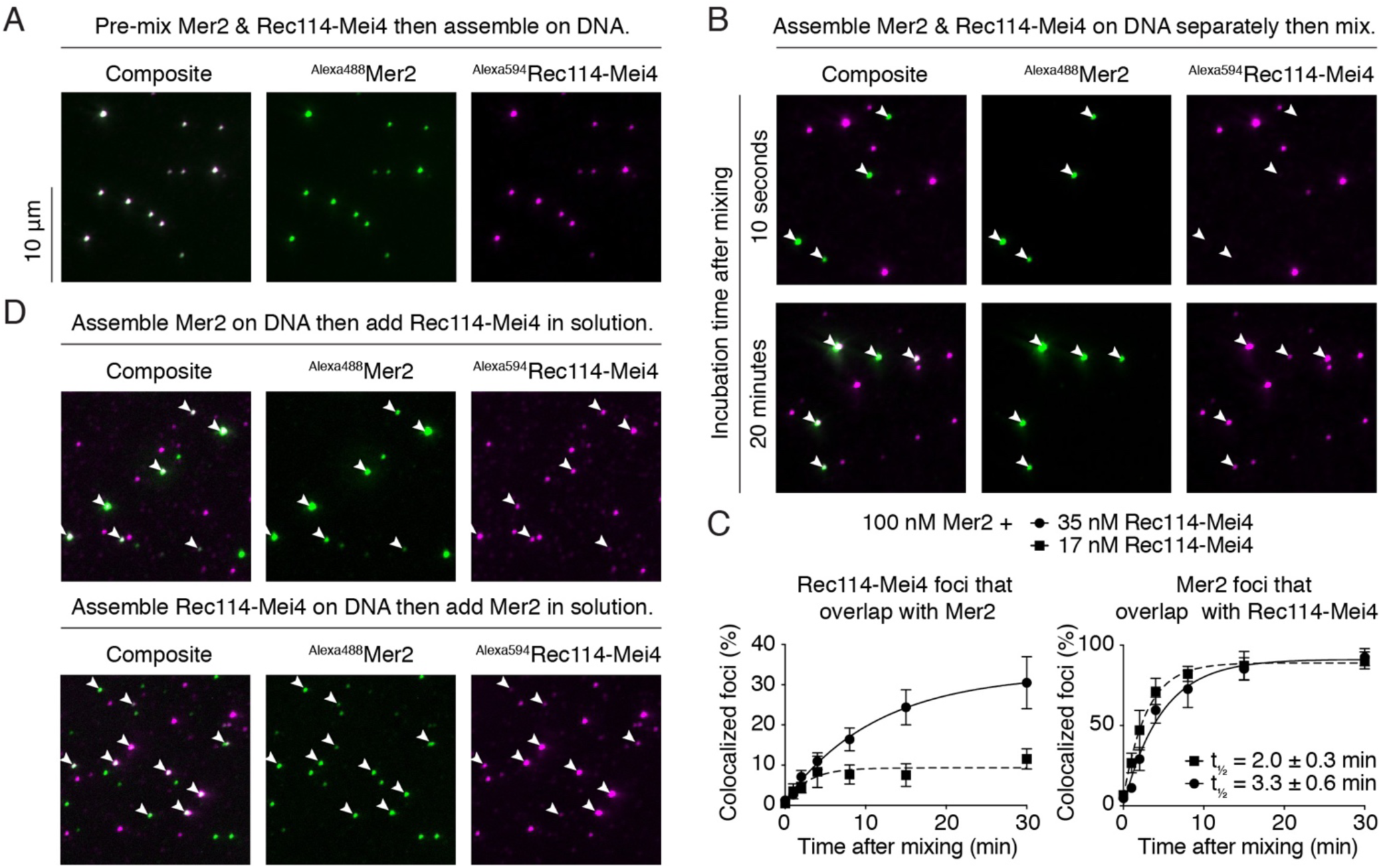
Assembly of tripartite Rec114–Mei4–Mer2 nucleoprotein condensates. A. Fluorescently labeled Rec114–Mei4 and Mer2 were mixed prior to DNA-driven condensation and imaged by fluorescence microscopy. Condensates were assembled for 30 minutes in the presence of 5.6 nM pUC19, 17 nM ^Alexa594^Rec114–Mei4 and 100 nM ^Alexa488^Mer2. B. Rec114–Mei4 and Mer2 nucleoprotein condensates were assembled separately for 10 minutes then mixed. After mixing, reactions contained 5.6 nM pUC19, 8.5 nM^Alexa594^Rec114–Mei4 and 50 nM ^Alexa488^Mer2. Samples were dropped on a microscope slide 10 seconds (top) or 20 minutes (bottom) after mixing. White arrowheads indicate positions of Mer2 condensates. C. Colocalization of Rec114–Mei4 and Mer2 in a condensate mixing time course. The time to achieve 50% of Mer2 foci overlapping with Rec114–Mei4 is indicated (t_½_). Lines are one-phase association models fit to the data. Error bars show mean ± SD from 8–10 fields of view. D. Recruitment of soluble Rec114–Mei4 (top) or Mer2 (bottom) into preassembled condensates of Mer2 (top) or Rec114–Mei4 (bottom). White arrowheads point to examples of the preassembled condensates.

Next, we addressed whether preassembled Rec114–Mei4 and Mer2 condensates have affinity for one another. When Rec114–Mei4 and Mer2 were condensed independently then mixed and the reactions plated immediately after mixing, no overlap was seen between the Rec114–Mei4 and Mer2 condensates, attributable to immobilization of the DNA on the glass surface (**Figure 5B**, top). In contrast, when reactions were incubated for 20 minutes prior to plating, all of the Mer2 condensates overlapped with a Rec114–Mei4 focus (**Figure 5B**, bottom). In principle, mixed foci could assemble either by fusion or by reassembly of foci via intermediate soluble protein. Since plating immobilizes DNA to the slide but not the protein, fusion should be inhibited by plating, but not dissociation/reassociation. Therefore, the lack of overlap in samples that were plated immediately rules out the scenario where mixed foci assemble via soluble intermediates.

To further test this and quantify the kinetics of focus fusion, we performed a time course experiment with two different concentrations of Rec114–Mei4 (17 and 35 nM) (**Figure 5C**). As shown in **Supplementary Figure 3A**, a higher concentration of Rec114–Mei4 yields a lower number of foci. If the fraction of joint clusters is driven by contact probability, the rate of overlap would be expected to be higher with the lower concentration of Rec114–Mei4. This was indeed the case: the rate at which Rec114–Mei4 was detected within Mer2 foci was higher with 17 nM (t_½_ = 2.0 ± 0.3 min) than with 35 nM (t_½_ = 3.3 ± 0.6 min) Rec114–Mei4 (**Figure 5C**, right panel).

Finally, we addressed whether soluble Rec114–Mei4 can be recruited into Mer2 condensates and vice versa. Here, Rec114–Mei4 or Mer2 condensates were assembled, then the partner subunit was added in solution, upon which the reactions were plated immediately to prevent subsequent focus fusion (**Figure 5D**). Preassembled Rec114– Mei4 foci contained signal from Mer2 and vice versa, showing that condensates provide nucleation sites for the partner complexes.

### Rec114–Mei4–Mer2 condensates recruit the Spo11 core complex

We sought to test the hypothesis that Rec114–Mei4–Mer2 condensates recruit Spo11 with its interacting partners Ski8, Rec102 and Rec104 (hereafter the “core complex”). To achieve this, we purified fluorescently labeled core complexes, bound the complex to DNA, then mixed the solution with preassembled Rec114–Mei4–Mer2 condensates. Consistent with our hypothesis, the core complex revealed punctate fluorescent signal that overlapped with Rec114–Mei4–Mer2 foci (**Figure 6A**). To evaluate specificity, we titrated Rec114–Mei4 and Mer2 and quantified core complex recruitment. Recruitment of the core complex to Rec114–Mei4 foci was dependent on Mer2 (**Figure 6B, D**). Surprisingly, when the reactions were performed with 100 nM Mer2, the core complex could be recruited to Mer2 foci independently of Rec114–Mei4 (**Figure 6C**). However, when Mer2 concentration was decreased to 25 nM, core complex recruitment became largely dependent on Rec114–Mei4 (**Figure 6C, E**).

**Figure 6:**
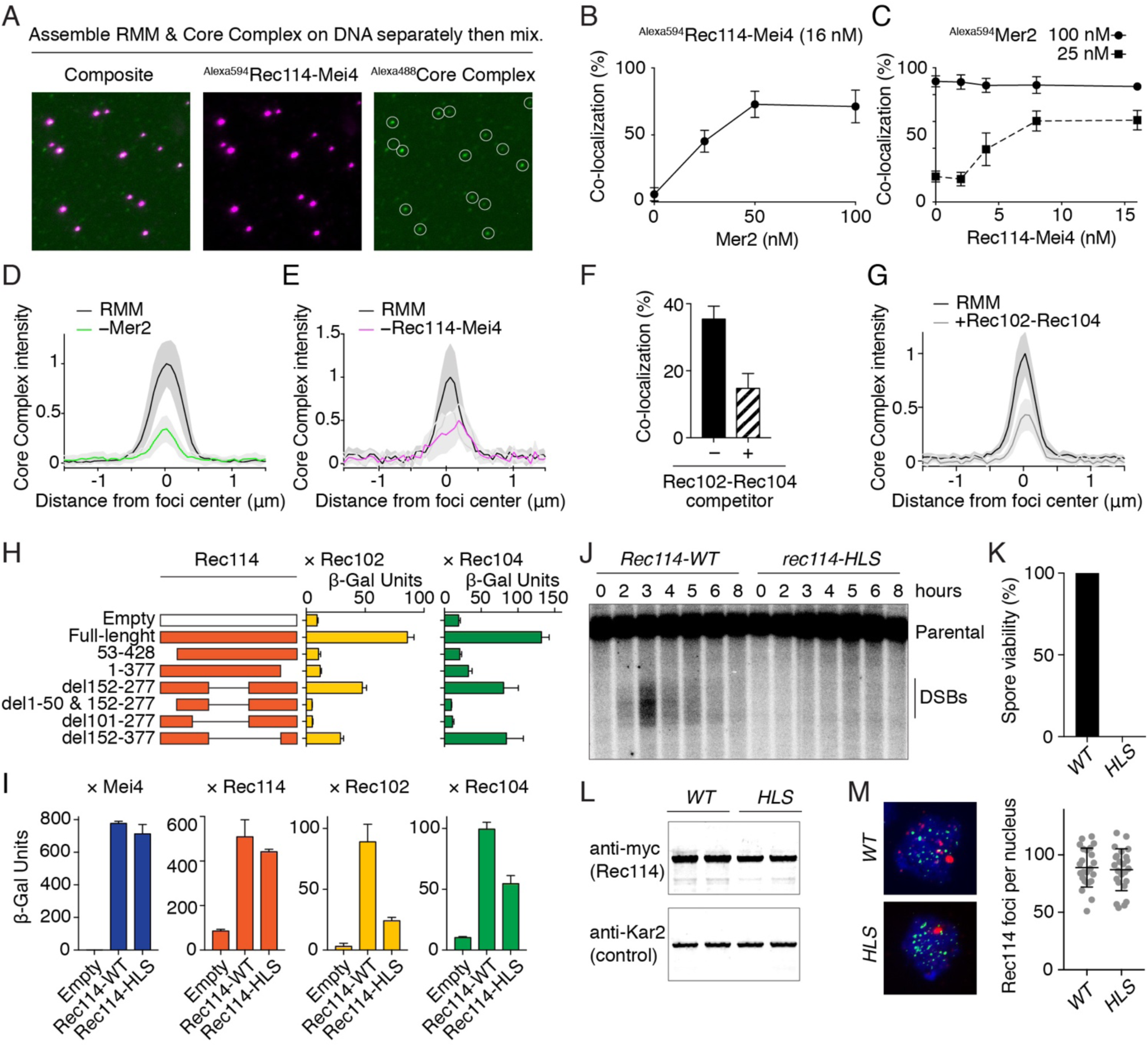
Rec114–Mei4–Mer2 condensates recruit the Spo11 core complex. A. Incorporation of Alexa488-labeled core complexes into Alexa594-labeled Rec114–Mei4–Mer2 condensates. B. Quantification of the fraction of Rec114–Mei4 foci that contain detectable core complex signal at the indicated concentration of Mer2. Error bars show mean ± SD from 8–10 fields of view. C. Quantification of the fraction of Mer2 foci that contain detectable core complex signal at the indicated concentration of Rec114–Mei4. Experiments were performed with either 25 nM or 100 nM Mer2. Error bars show mean ± SD from 8–10 fields of view. D. Quantification of core complex signal within Rec114–Mei4 foci in the presence (100 nM) or absence of Mer2. The average intensity within 20 foci is plotted for each reaction. Shaded areas represent 95% confidence intervals. E. Quantification of core complex signal within Mer2 foci in the presence (16 nM) or absence of Rec114–Mei4. Reactions contained 25 nM Mer2. The average intensity within 20 foci is plotted for each reaction. Shaded areas represent 95% confidence intervals. F. Effect of including 100 nM ^MBP^Rec102–Rec104^HisFlag^ competitor on the recruitment of the core complex to RMM condensates (16 nM Rec114–Mei4, 25 nM Mer2). The fraction of Rec114–Mei4–Mer2 foci that contain detectable core complex signal is plotted. Error bars show mean ± SD from 8–10 fields of view. G. Intensity of core complex signal within Rec114–Mei4–Mer2 condensates in the absence or presence of Rec102–Rec104 competitor. The average core complex intensity within 20 foci is plotted for each reaction. Shaded areas represent 95% confidence intervals. H. Mapping regions of Rec114 required for interaction with Rec102 or Rec104 by Y2H analysis. β-galactosidase units are measured for the interaction between truncated variants of Gal4AD-Rec114 and LexA-Rec102 or LexA-Rec104. Error bars represent SD from four replicates. I. Quantification of Y2H interaction between Gal4AD-Rec114 wild type or H39A/L40A/S41A (HLS) mutant and LexA-Mei4, LexA-Rec114, LexA-Rec102, or LexA-Rec104. Error bars represent SD from four replicates. J. Southern blot analysis of meiotic DSB formation at the *CCT6* hotspot. K. Spore viability of Rec114-WT and HLS mutant strains. L. Western-blot analysis of meiotic protein extracts from myc-tagged Rec114-WT and HLS mutant strains. Samples from two biological replicates are shown. M. Immunofluorescence microscopy of meiotic chromosome spreads with myc-tagged Rec114-WT and HLS mutant strains. Green, anti-myc (Rec114); red, anti-Zip1; blue, DAPI. Quantification of the number of Rec114 foci per leptotene or early zygotene cell is plotted. Error bars represent SD.

Rec114 interacts with the core complex subunits Rec102 and Rec104 in Y2H assays (Arora et al., 2004; Maleki et al., 2007). We therefore asked whether an excess of Rec102–Rec104 subcomplexes would compete with the full core complex for recruitment within the condensates. This was indeed the case (**Figure 6F, G**).

We used a Y2H assay to map the core complex interacting domain of Rec114 by truncation analysis (**Figure 6H**). Deleting ∼50 amino acids from either the N or C termini of Rec114 was sufficient to abolish the Y2H interaction with both Rec102 and Rec104, but a deletion of amino acids 152-377 (predicted to be disordered, see **Figure 1H**) maintained the protein-protein interaction. We mutated conserved residues within the N-terminal PH domain of Rec114 in search for mutations that specifically compromised the interaction with Rec102 and Rec104, but did not affect the interaction with Mei4 and wild-type Rec114, and identified a mutation (HLS, H39A/L40A/S41A) that fulfilled these criteria (**Figure 6I**).

To address whether the HLS mutation affects Rec114 function, we generated a yeast strain that carried the mutation and quantified meiotic DSB formation and spore viability. Southern blot analysis at the *CCT6* hotspot showed that the *rec114-HLS* mutant was defective for DSB formation (**Figure 6J**), which lead to inviable spores (**Figure 6K**). Nevertheless, western-blot analysis and immunofluorescence of meiotic chromosome spreads using a myc-tagged Rec114 version showed that the HLS mutation did not affect protein expression (**Figure 6L**) or the formation of chromatin-associated foci (**Figure 6M**).

Together, these data are consistent with the idea that the core complex is recruited to Rec114–Mei4–Mer2 condensates, probably through at least two sets of interactions: one that depends on Mer2 and another that likely involves contacts between the N-terminal PH domain of Rec114 and both the Rec102 and Rec104 subunits of the core complex.

## Discussion

We have shown that Rec114–Mei4 and Mer2 form separate subcomplexes *in vitro* that each bind DNA with a remarkable degree of cooperativity and assemble micrometer-scale nucleoprotein super-complexes. The DNA-dependent clusters are reminiscent of biomolecular condensates that control a variety of processes (Li et al., 2012; Su et al., 2016; Wheeler et al., 2016; Banani et al., 2017; Boulay et al., 2017; Boeynaems et al., 2018). Specifically, Rec114–Mei4 and Mer2 nucleoprotein assemblies are reversible, promoted by molecular crowding, and can fuse. They appear to depend on multivalent interactions between the negatively charged DNA backbone and positively charged residues within the proteins. We identified residues in Rec114 and Mer2 that are important for DNA binding and condensation *in vitro* and formation of chromatin-associated Rec114 and Mer2 foci and meiotic DSB formation *in vivo*. These direct relationships between *in vitro* and *in vivo* activities suggest that DNA-driven condensation is an important aspect of the biological function of Rec114, Mei4 and Mer2.

Previous work resulted in Rec114, Mei4, and Mer2 being loosely grouped together as a functionally related subset of the DSB-essential proteins, but it was never clear precisely how they interacted with one another (Lam and Keeney, 2015). Mer2 foci are independent of Rec114 and Mei4, but chromatin association of Rec114 and Mei4 appears to largely depend on each other and on Mer2 (Li et al., 2006; Maleki et al., 2007; Panizza et al., 2011). These different dependencies are also the case in mice (Stanzione et al., 2016; Kumar et al., 2018; Acquaviva et al., 2019). Our results provide a concrete molecular framework for understanding these previously unclear relationships: heterotrimeric Rec114–Mei4 complexes and Mer2 homotetramers are biochemically distinct entities that each bind cooperatively to DNA, but they collaborate in DSB formation in the context of mixed, DNA-dependent condensates.

Chromatin-associated Rec114, Mei4 and Mer2 foci localize to chromosome axes in *S. cerevisiae, S. pombe* (Rec7, Rec24, and Rec15) and mice (REC114, MEI4 and IHO1), and probably also in other organisms including plants (PHS1, PRD2 and PRD3/PAIR1) (Pawlowski et al., 2004; Henderson et al., 2006; Li et al., 2006; Lorenz et al., 2006; Maleki et al., 2007; De Muyt et al., 2009; Kumar et al., 2010; Bonfils et al., 2011; Panizza et al., 2011; Miyoshi et al., 2012; Kumar et al., 2018). We therefore consider that the DNA-driven condensation we uncovered here is likely to be a fundamental, evolutionarily conserved property of these proteins. The synaptonemal complex is another example of how phase separation is proposed to govern spatial patterning of meiotic recombination (Rog et al., 2017).

### Phase separation and the molecular basis of Rec114–Mei4–Mer2 assemblies

What is the mechanistic basis for the clustering behavior of Rec114–Mei4 and Mer2? Our experiments reveal close similarities between DNA-driven condensation of Rec114–Mei4 and Mer2 and systems that undergo phase separation. Two types of mechanisms have been proposed to explain the self-assembly of chromatin subcompartments by phase separation: one involves a multivalent chromatin binder, the second involves a self-associating chromatin binder. Both types of interactions can induce phase separation of nuclear bodies around the protein binding sites (Erdel and Rippe, 2018).

The fact that both Rec114–Mei4 and Mer2 are multimers suggest that they may constitute multivalent binders that crosslink the DNA scaffold. Such interactions are predicted to fall apart upon removal of the DNA, which we observed when challenging the condensates by DNase I treatment. In contrast, protein droplets resulting from self-associating chromatin binders are predicted to persist after DNA removal (Erdel and Rippe, 2018). Nevertheless, multivalent DNA binding alone cannot explain the highly cooperative assembly of Rec114–Mei4 and Mer2 that restricts itself to just a subset of available DNA molecules, so protein self-association must also be invoked. These self interactions are presumably too weak to persist upon DNA removal.

Multivalent interactions on the DNA scaffold lead to amorphous, stochastic assemblies with no defined higher-order structure. We expect protein diffusion within and in-and-out of the condensates to be dynamic, but the extent to which this is true *in vivo* is unclear. *In vitro*, in-and-out diffusion is highly influenced by experimental conditions and is reduced by a crowding agent. Fusion is a major factor for focus growth *in vitro*, but it is likely that chromosomal drag limits the fusion of condensates *in vivo*.

Depending on the strength of the interactions, phase-separated systems can exist as liquid, gel-like or even solid forms, and transitions from liquid to solid may occur spontaneously (Lin et al., 2015; Patel et al., 2015; Xiang et al., 2015; Banani et al., 2017). We found that the reversibility of the Rec114–Mei4–Mer2 condensates decreases over time, influenced by molecular crowding, potentially consistent with the progressive transitions to gel-like or solid states.

### A model for the assembly of the meiotic break machinery by DNA-driven Rec114–Mei4–Mer2 condensation

A longstanding question has been how the different DSB proteins promote Spo11 activity. Based on their axis-association and the physical link between Mer2, Spp1 and H3K4 trimethylation, which marks regions of preferential DSB activity within chromatin loops, Rec114, Mei4 and Mer2 have been proposed to form part of the tethered loop-axis structure that is thought to assemble in preparation for Spo11-mediated DNA cleavage (Arora et al., 2004; Henderson et al., 2006; Li et al., 2006; Maleki et al., 2007; Panizza et al., 2011; Acquaviva et al., 2013; Sommermeyer et al., 2013; Cooper et al., 2016). Our data adds to this model by highlighting the organizing role of Rec114–Mei4 and Mer2 in the assembly of punctate clusters. We propose that it is these clusters that provide the structural assembly for recruitment of Spo11 and other regulatory components, and that it is in the context of these structures that the tethered loop-axis configuration is eventually adopted (**Figure 7A**).

**Figure 7:**
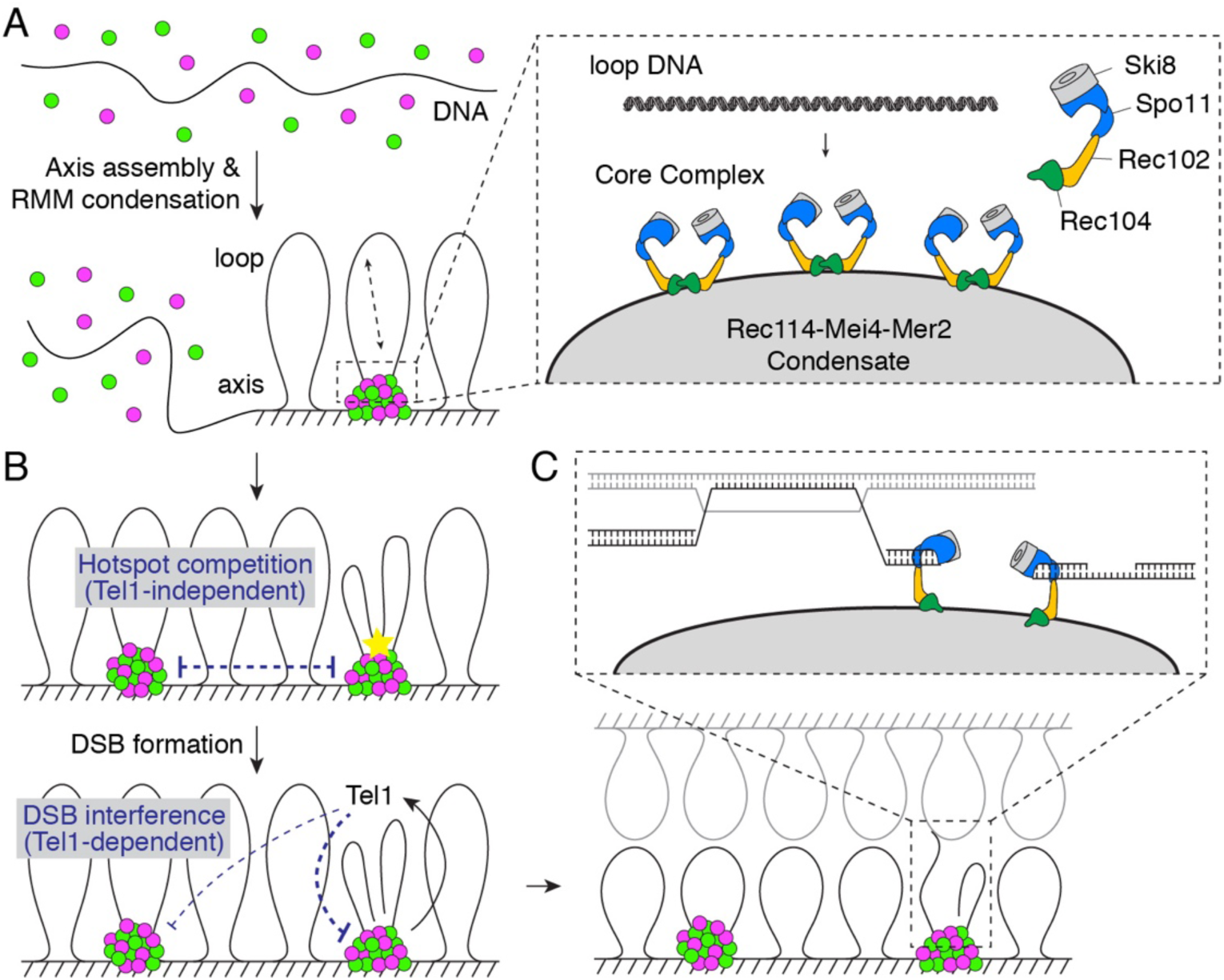
A condensate model for assembly of the meiotic DSB machinery and implications for the control of DSB formation and repair. A. Assembly of the DSB machinery. (Left) Rec114–Mei4 and Mer2 complexes bind DNA in a highly cooperative manner to form large mixed nucleoprotein condensates. (Right) These condensates provide a platform to recruit the core complex through interactions that involve the N-terminal domain of Rec114 and the Rec102–Rec104 components of the core complex. Multiple Spo11 complexes are recruited and may engage an incoming DNA loop simultaneously. The molecular arrangement of the core complex proteins is based on (Claeys Bouuaert *et al.*, in preparation). B. Hotspot competition and DSB interference. Competition arises prior to DSB formation as a consequence of the partitioning of RMM proteins into condensates. DSB interference is implemented through local inhibition of further DSB formation by DSB-activated Tel1. Inhibition could work on the same cluster that generated the activation DSB as well as on nearby clusters in cis. C. The coherence provided by the condensates may serve functions during repair, including the maintenance of a physical connection between the DNA ends that involves end-capping by condensate-embedded core complexes.

Hyperstoichiometric condensates would be expected to recruit multiple core complexes, which is supported by our *in vitro* data and the detection of Spo11 foci *in vivo* (Prieler et al., 2005). Each cluster may thus be expected to be capable of forming multiple DSBs. Indeed, closely spaced pairs of DSBs on the same chromatid (double cuts) occur at a much higher frequency than expected by chance, estimated to account for ∼10% of total Spo11 activity in wild-type cells (Johnson et al., 2019). The distance between pairs of DSBs ranges from about 30 to >100 bp with a 10-bp periodicity, corresponding to the pitch of the double helix. This periodicity can be explained if two Spo11 complexes engage DNA in the same orientation, which in turn can be explained if the complexes are constrained on a surface (Johnson et al., 2019). In the light of our findings, we propose that Rec114–Mei4–Mer2 condensates provide platforms that recruit and display co-oriented arrays of Spo11 complexes that then capture and break loop DNA (**Figure 7A**). Whether the core complex is mostly at the surface of the condensates or is only active when exposed on the surface is unclear.

### Hotspot competition and DSB interference

Our model has implications for numerical and spatial control of DSB patterning. It has long been appreciated that the presence of a strong hotspot can reduce DSB activity of a neighboring hotspot. This phenomenon—hotspot competition—is a population-average effect (Wu and Lichten, 1995; Xu and Kleckner, 1995; Fan et al., 1997; Jessop et al., 2005; Robine et al., 2007; Mohibullah and Keeney, 2017). Another process—DSB interference—is observed when cleavage of individual DNA molecules within a cell is considered: the presence of a DSB decreases the chances of another DSB occurring nearby, at distances up to a hundred kilobases (Garcia et al., 2015).

Hotspot competition and DSB interference are genetically separable in *S. cerevisiae* because DSB interference depends on the DNA-damage response kinase Tel1 (Garcia et al., 2015), while hotspot competition does not (Mohibullah and Keeney, 2017). Tel1 governs a *cis*-acting DSB-dependent negative feedback loop (Lange et al., 2011; Zhang et al., 2011). In a *tel1* mutant, not only is DSB interference eliminated, but negative interference is detected over distances on the ∼10 kb scale, meaning that coordinated cutting of the same chromatid at adjacent hotspots is observed at high frequency (Garcia et al., 2015).

Our model for locally coordinated break formation through the recruitment of multiple Spo11 complexes within Rec114–Mei4–Mer2 condensates provides a molecular explanation for these behaviors. Hotspot competition could be implemented prior to break formation and can be explained if hotspots compete for poorly diffusing and locally limiting factors (Wu and Lichten, 1995). Indeed, Rec114–Mei4–Mer2 have been proposed to constitute this factor based on the observation that they associate to the chromosome axes, which may therefore limit their diffusion within the nucleus, hence their availability (Panizza et al., 2011). The highly cooperative DNA-driven condensation described here provides a molecular basis to understand how this might work, because the nucleation of a condensate would cause a local depletion of Rec114–Mei4 and Mer2 proteins, reducing the probability of another nucleation event (**Figure 7B**). After a DSB is made, Tel1 is envisioned to act both within and between adjacent condensates to suppress additional DSBs nearby (**Figure 7B**).

Alternatively, it has been proposed that hotspot competition and DSB interference could reflect association of a cluster of several chromatin loops with a limited-catalytic-capacity DSB-forming assembly (Garcia et al., 2015; Cooper et al., 2016). In this view, loops within a cluster compete with one another for access to the DSB machinery. Hotspot clustering has also been proposed as a means to control DSB patterning in *S. pombe* (Fowler et al., 2018). Once activated, the machinery has potential to make multiple cuts, but this is suppressed by Tel1 after the first DSB is made. How such loop clustering might occur and how it might be integrated with the DSB machinery have been a matter of conjecture, and mechanisms behind speculated roles of Rec114, Mei4, and Mer2 were likewise unclear (Cooper et al., 2016). The macromolecular condensates we document could readily account for all of these properties.

### Macromolecular condensates as a platform to integrate DSB formation with repair

An intriguing possibility is that forming DSBs in the context of a superstructure may facilitate control of subsequent repair. The condensates themselves, being axis-associated, could hold the bases of the broken DNA loop, providing a coherent unit that would prevent the diffusion of the DNA ends from each other (**Figure 7B**, bottom panel). In addition, one or both DSB ends might remain embedded within the condensate via the persistence of Spo11-oligos that might cap the ends after resection. End-capping by Spo11 (Neale et al., 2005) has recently received support from patterns of recombination intermediates detected in mice (Paiano et al., 2019; Yamada et al., 2019), and is consistent with the observation that Spo11 binds tightly to DNA ends *in vitro*, even in the absence of a covalent link (Claeys Bouuaert *et al.*, in preparation).

Following DSB formation, one might imagine that the condensates hand over or evolve into recombination nodules where DSB repair takes place. In *Sordaria macrospora*, the Mer2 ortholog Asy2 re-localizes from the axis to the central region of the synaptonemal complex as the latter assembles during zygonema (Tesse et al., 2017). Similar movements are also seen for recombination proteins Mer3 and Msh4 (Storlazzi et al., 2010; Espagne et al., 2011), but Mer2 re-localization to the axis is independent of Mer3 and Msh4, suggesting that Mer2 may have a role in transporting recombination complexes to the nascent synaptonemal complex. We propose that this transport takes advantage of the coherence provided by the DNA-dependent condensates.

In summary, our results suggest a model whereby DNA-driven condensation of Rec114–Mei4 and Mer2 organize punctate compartments along meiotic chromosomes that provide the structural context to facilitate controlled break formation and perhaps downstream repair. This molecular framework explains the phenomena of hotspot competition and the occurrence and periodicity pattern of closely-spaced DSBs, and provides insights into the relationships between DSB formation and the loop-axis structure of chromosomes.

## Materials and Methods

### Preparation of expression vectors

Oligonucleotides (oligos) used in this study were purchased from Integrated DNA Technologies. The sequence of the oligos is listed in **Supplementary Table S2**. Plasmids are listed in **Supplementary Table S3**.

Separate exons of *S. cerevisiae REC114, MEI4* and *MER2* were amplified from genomic DNA of the SK1 strain and assembled by in-fusion cloning to yield intron-less pFastbac1-derived expression vectors pCCB649, pCCB652 and pCCB681, respectively. Primers for *REC114* were: cb906 and cb907 (exon 1), and cb908 and cb909 (exon 2). Primers for *MEI4* were: cb910 and cb911 (exon 1), and cb912 and 913 (exon 2). Primers for *MER2* were: cb978 and cb979 (exon 1), and cb980 and cb981 (exon 2). The genes were subcloned into pFastBac-HTb-Flag to generate N-terminally HisFlag-tagged expression vectors for ^HisFlag^Rec114 (pCCB650), Mei4 (pCCB653) and Mer2 (pCCB682). MBP was subcloned into the untagged vectors, to yield expression vectors for ^MBP^Rec114 (pCCB651) and ^MBP^Mei4 (pCCB654), ^MBP^Mer2 (pCCB683).

To generate untagged Rec114–Mei4, the cleavage sequence for the TEV protease was introduced between the affinity tag and the sequence coding for Rec114 and Mei4 by inverse PCR and self-ligation using templates pCCB650 (primers cb1283 and cb1284), and pCCB654 (primers cb1287 and cb1288), to yield vectors pCCB789 (^HisFlag-TEV^Rec114) and pCCB791 (^MBP-TEV^Mei4), respectively. The mScarlet fluorophore was amplified from a synthetic gene codon-optimized for mammalian expression (gift from Soonjoung Kim, MSKCC) with primers cb1279 and cb1280 and cloned into the BamHI site of pCCB650 to yield pCCB786 (^HisFlag-mScarlet^Rec114). A TEV cleavage site was further introduced between the affinity tag and the fluorophore by inverse PCR and self-ligation using template pCCB786 and primers cb1285 and cb1286 to yield pCCB790 (^HisFlag-TEV-mScarlet^Rec114). The Rec114-R395A/K396A/K399A/K400A (4KR) mutant was generated by inverse-PCR and self-ligation of pCCB789 and pCCB790 with primers cb1332 and cb1334 to yield pCCB848 (^HisFlag-TEV^Rec114-4KR), pCCB849 (^HisFlag-TEV-mScarlet^Rec114-4KR).

To generate a vector for Mer2 expression in *E. coli, MER2* was amplified from pCCB681 using primers cb1161 and cb1162 and cloned into the BamHI site of pSMT3 to yield pCCB750 (^SUMO^Mer2). eGFP-tagged Mer2 was generated by PCR amplification of eGFP using primers cb1259 and cb1260 and in-fusion cloning in the BamHI site of pCCB750 to yield pCCB777 (^SUMO-eGFP^Mer2). The Mer2-K265A/R266A/R267A/R268A (KRRR) mutation was generated by QuikChange mutagenesis using primers cb1186 and cb1187 pf pCCB750 and pCCB777 to yield pCCB779 (^SUMO^Mer2-KRRR) and pCCB783 (^SUMO-eGFP^Mer2-KRRR), respectively.

Full-length Rec114 and Mei4 were amplified from pCCB649 and pCCB650 using primers sp16 and sp17, and sp25 and sp26, respectively and cloned into the pETDuet-1 vector by in-fusion cloning to yield pSP34. A SUMO tag was introduced at the N-terminus of Rec114 by PCR amplification of the pSMT3 vector with primers cb1172 and cb1180 and In-Fusion cloning within the Nco1 and BamHI fragment of pSP34 to yield pSP53. Truncations were obtained from this construct by inverse PCR and self-ligation.

### Expression and purification of recombinant proteins

Viruses were produced by a Bac-to-Bac Baculovirus Expression System (Invitrogen) following the manufacturer’s instructions. 2 × 10^9^ *Spodoptera frugiperda* Sf9 cells were infected with combinations of viruses at a multiplicity of infection (MOI) of 2.5 each. Expression of ^HisFlag^Rec114-^MBP^Mei4 used viruses generated from pCCB650 and pCCB654, untagged Rec114–Mei4 used viruses generated from pCCB789 and pCCB791, and fluorescently tagged ^mScarlet^Rec114–Mei4 used viruses generated from pCCB790 and pCCB791. After 62 h infection, cells were harvested, washed with phosphate buffer saline (PBS), frozen in dry ice and kept at −80 °C until use. All the purification steps were carried out at 0–4 °C. Cell pellets were resuspended in 4 volumes of lysis buffer (25 mM HEPES-NaOH pH 7.5, 500 mM NaCl, 0.1 mM DTT, 20 mM imidazole, 1x Complete protease inhibitor tablet (Roche) and 0.1 mM phenylmethanesulfonyl fluoride (PMSF)). Cells were lysed by sonication and centrifuged at 43,000 g for 30 min. The cleared extract was loaded onto 1 ml pre-equilibrated NiNTA resin (Qiagen). The column was washed extensively with Nickel buffer (25 mM HEPES-NaOH pH 7.5, 500 mM NaCl, 10% glycerol, 0.1 mM DTT, 20 mM imidazole, 0.1 mM PMSF). The tagged complexes were then eluted in Nickel buffer containing 250 mM imidazole. The complexes were further purified on amylose resin (NEB). Fractions containing protein were pooled and diluted in 3 volumes of Amylose buffer (25 mM HEPES-NaOH pH 7.5, 500 mM NaCl, 10% glycerol, 2 mM DTT, 5 mM EDTA). Next, the complexes were bound to 1 ml of the Amylose resin in a poly-prep chromatography column (Bio-Rad) and the resin was washed extensively. Complexes were eluted from amylose resin with buffer containing 10 mM maltose. Fractions containing protein were pooled and loaded on a Superdex 200 column preequilibrated with Amylose buffer. For untagged or mScarlet-tagged complexes, samples were treated with an excess of TEV protease prior to gel filtration. For fluorescently labeled complexes, labeling was performed using Alexa Fluor 594 (Invitrogen #A10239), which has a succinimidyl ester moiety that reacts with primary amines. After 1 hour conjugation at room temperature, complexes were purified by gel filtration. Fractions containing protein were concentrated in 50 kDa cutoff Amicon centrifugal filters (Millipore). Aliquots were frozen in dry ice and stored at −80 °C.

For expression of recombinant proteins in *E. coli*, expression vectors were transformed in BL21 DE3 cells and plated on LB plates containing the appropriate antibiotic. Cells were then cultured in liquid medium at 37 °C to OD_600_ = 0.6. For Mer2 proteins and variants, expression was carried out at 30 °C for 3 hours with 1 mM IPTG. For Rec114–Mei4 truncations, expression was carried out at 16 °C overnight with 0.2 mM IPTG. Cells were resuspended in Nickel buffer (25 mM HEPES-NaOH pH 7.5, 500 mM NaCl, 10% glycerol, 0.1 mM DTT, 20 mM imidazole, 0.1 mM PMSF) and frozen dropwise in liquid nitrogen and kept at −80 °C until use. All the purification steps were carried out at 0–4 °C. Cells were lysed using a French press and centrifuged at 43,000 g for 30 min. The cleared extract was loaded onto 1 ml pre-equilibrated NiNTA resin (Qiagen). The column was washed extensively with Nickel buffer then eluted in buffer containing 250 mM imidazole. The 6His-SUMO tag was cleaved with Ulp1 during overnight dialysis in gel filtration buffer (25 mM HEPES-NaOH pH 7.5, 300 mM NaCl, 10% glycerol, 40 mM imidazole, 1 mM DTT, 5 mM EDTA). The sample was then loaded on a second Nickel column to remove 6His-SUMO and Ulp1. The flow-through was then loaded on a Superdex 200 column preequilibrated with gel filtration buffer. For Mer2 complexes labeled with Alexa Fluor 488 (Invitrogen #A10235), fluorophore conjugation was performed at room temperature for 1 hour prior to gel filtration. Alexa Fluor 488 has a tetrafluorophenyl ester moiety that reacts with primary amines. After gel filtration, fractions containing protein were concentrated in 10 kDa cutoff Amicon centrifugal filters (Millipore). Aliquots were frozen in dry ice and stored at −80 °C.

### SEC-MALS

Light scattering data in **Figure 1C, H** were collected using a Superdex 200, 10/300, HR Size Exclusion Chromatography (SEC) column (GE Healthcare, Piscataway, NJ), connected to High Performance Liquid Chromatography System (HPLC), Agilent 1200, (Agilent Technologies, Wilmington, DE) equipped with an autosampler. The elution from SEC was monitored by a photodiode array (PDA) UV/VIS detector (Agilent Technologies, Wilmington, DE), differential refractometer (OPTI-Lab rEx Wyatt Corp., Santa Barbara, CA), static and dynamic, multiangle laser light scattering (LS) detector (HELEOS II with QELS capability, Wyatt Corp., Santa Barbara, CA). The SEC-UV/LS/RI system was equilibrated in buffer 25 mM Hepes pH 7.5, 500 mM NaCl, 10 % glycerol, 2 mM EDTA at the flow rate of 0.5 ml/min or 1.0 ml/min. Two software packages were used for data collection and analysis: the Chemstation software (Agilent Technologies, Wilmington, DE) controlled the HPLC operation and data collection from the multi-wavelength UV/VIS detector, while the ASTRA software (Wyatt Corp., Santa Barbara, CA) collected data from the refractive index detector, the light scattering detectors, and recorded the UV trace at 280 nm sent from the PDA detector. The weight average molecular masses were determined across the entire elution profile in intervals of 1 sec from static LS measurement using ASTRA software.

All other SEC-MALS experiments were performed by an Äkta-MALS system. Proteins (500 µl) were loaded on Superdex 75 10/300 GL or Superdex 200 10/300 GL columns (GE Healthcare) and eluted with buffer 20 mM Tris pH 7.5, 300 mM NaCl, 2 mM DTT at a flow rate of 0.3 ml/min. The light scattering was monitored by a miniDAWN TREOS system (Wyatt Technologies) and concentration was measured by an Optilab T-rEX differential refractometer (Wyatt Technologies).

### Crosslinking – mass spectrometry

For crosslinking, ∼20–50 µg of ^HisFlag^Rec114–^MBP^Mei4 or ^HisFlag^Mer2 complexes were incubated in 50–100 µl reactions in the presence of 2 mM disuccinimidyl suberate (DSS) in buffer containing 25 mM HEPES-NaOH pH 7.5, 500 mM NaCl, 10% glycerol, 2 mM DTT, 5 mM EDTA. After 10 minutes crosslinking at 30 °C, reactions were quenched with 100 mM Tris-HCl pH 7.5. Crosslinked proteins were separated by SDS-polyacrylamide gel electrophoresis and stained with SimplyBlue (Invitrogen). Protein bands were excised and digested *in situ* with trypsin as described (Sebastiaan Winkler et al., 2002). The tryptic peptides were purified using a 2-µl bed volume of Poros 50 R2 (Applied Biosystems) reverse-phase beads packed in Eppendorf gel-loading tips (Erdjument-Bromage et al., 1998). The digested peptides were diluted in 0.1% formic acid, and each sample was analyzed separately by microcapillary LC with tandem MS by using the NanoAcquity system (Waters) with a 100 µm inner diameter × 10 cm length C18 column (1.7 µm BEH130; Waters) configured with a 180 µm × 2 cm trap column coupled to a Q-Exactive Plus mass spectrometer (Thermo Fisher Scientific). A proxeon nanoelectrospray source set at 1800 V and a 75 µm (with 10 µm orifice) fused silica nano-electrospray needle (New Objective, Woburn, MA) was used to complete the interface. 1 µl of sample was loaded onto the trap column, washed with 3x loop volume of buffer A (0.1% formic acid) and the flow was reversed through the trap column and the peptides eluted with a 1-50% acetonitrile (with 0.1% formic acid) gradient over 50 min at a flow rate of 300 nl/min over the analytical column. The QE Plus was operated in automatic, data-dependent MS/MS acquisition mode with one MS full scan (370–1700 *m*/*z*) at 70,000 mass resolution and up to ten concurrent MS/MS scans for the ten most intense peaks selected from each survey scan. Survey scans were acquired in profile mode and MS/MS scans were acquired in centroid mode at 17500 resolution and isolation window of 1.5 amu. AGC was set to 1 × 10^6^ for MS1 and 5 × 10^5^ and 100 ms maximum IT for MS2. Charge exclusion of 1, 2 and greater than 8 enabled with dynamic exclusion of 15 s. To analyze the cross-linked peptides we used pLink (Yang et al., 2012). The raw MS data was analyzed using pLink search with the following parameters: precursor mass tolerance 50 p.p.m., fragment mass tolerance 10 p.p.m., cross-linker DSS (cross-linking sites K and protein N terminus), xlink mass-shift 138.068, monolink mass-shift 156.079, fixed modification C 57.02146, variable modification oxidized methionine, deamidation N,Q, protein N-acetyl, peptide length minimum 4 amino acids and maximum 100 amino acids per chain, peptide mass minimum 400 and maximum 10,000 Da per chain, enzyme trypsin, two missed cleavage sites per chain (four per cross-link). The data were imported on the xiNET online tool to generate crosslinking maps (Combe et al., 2015). All identified crosslinks can be found in **Supplementary Table S1**.

To estimate the ratio of Rec114 and Mei4 by mass spectrometry, 10 µg of ^HisFlag^Rec114–^MBP^Mei4 were digested with trypsin, analyzed by tandem MS as described above, and spectral counts of the two proteins were compared, omitting the tags. Rec114 and Mei4 have similar lengths (428 and 408 amino acids, respectively), and similar numbers of K and R residues (56 and 66 respectively). The average and median trypic peptide length is 7.6 and 5 for Rec114, and 6.1 and 4 for Mei4. The .raw files were converted to .mgf and searched by Mascot (Matrix Science, version 2.6.100) using the Fasta formatted Swissprot reviewed database (downloaded July 5, 2017 from www.UniProt.org) and the Fasta formatted Rec114 and Mei4 sequence. The search parameters were as follows: (i) two missed cleavage tryptic sites were allowed; (ii) precursor ion mass tolerance 10 ppm; (iii) fragment ion mass tolerance 0.08 Da (monoisotopic); and (iv) fixed modification of carbamidomethyl of cysteine; (v) variable protein modifications were allowed for methionine oxidation, deamidation on NQ, protein N-terminal acetylation, and phospho STY. Scaffold (version Scaffold_4.8.4, Proteome Software Inc., Portland, OR) was used to validate MS/MS based peptide and protein identifications. Peptide identifications were accepted if they could be established at greater than 70% probability to achieve an FDR less than 1% by the Scaffold Local FDR algorithm. Protein identifications were accepted if they could be established at greater than 6% probability to achieve an FDR less than 1% and contained at least 2 identified peptides. Protein probabilities were assigned by the Protein Prophet algorithm (Nesvizhskii et al., 2003). Proteins that contained similar peptides and could not be differentiated based on MS/MS analysis alone were grouped to satisfy the principles of parsimony. Proteins sharing significant peptide evidence were grouped into clusters.

### AFM imaging

For AFM imaging of Rec114–Mei4 or Mer2 bound to plasmid DNA, protein complexes were diluted to the indicated concentration (12–50 nM) in the presence of 1 nM supercoiled pUC19 in 25 mM HEPES-NaOH pH 6.8, 5 mM MgCl_2_, 50 mM NaCl, 10% glycerol. Complexes were assembled at 30 °C for 30 minutes. A volume of 40 µl of the protein-DNA binding reaction was deposited onto freshly cleaved mica (SP1) for 2 minutes. The sample was rinsed with 10 ml ultrapure deionized water and the surface was dried using a stream of nitrogen. AFM images were captured using an Asylum Research MFP-3D-BIO (Oxford Instruments) microscope in tapping mode at room temperature. An Olympus AC240TS-R3 AFM probe with resonance frequencies of approximately 70 kHz and spring constant of approximately 1.7 N/m was used for imaging. Images were collected at a speed of 0.5–1 Hz with an image size of 2 µm at 2048 × 2048 pixel resolution.

### DNA substrates and gel shift assays

Short linear DNA substrates were generated by annealing complementary oligos (sequences listed in **Supplementary Table S2**). The substrates were the following (with oligo names in parentheses): dsDNA20 (cb939 & cb940), dsDNA40 (cb922 & cb935), dsDNA80 (cb95 & cb100). The 80 nt oligos were first purified on 10% polyacrylamide-urea gels. Oligos were subsequently mixed in equimolar concentrations (10 µM) in STE (100 mM NaCl, 10 mM Tris-HCl pH 8, 1 mM EDTA), heated and slowly cooled on a PCR thermocycler (98°C for 3 min, 75°C for 1 h, 65°C for 1 h, 37°C for 30 min, 25°C for 10 min). For radioactive labeling, 1/20^th^ of the annealed substrates were 5′-end-labeled with [γ-^32^P]-ATP (Perkin Elmer) and T4 polynucleotide kinase (New England Biolabs). Labeled and unlabeled substrates were purified by native polyacrylamide gel electrophoresis. Larger linear substrates were prepared by PCR amplification of a 9.6-kb template derived from pUC19 (pDR470). Substrates were as follows: 100 bp (cb343 & cb1339), 1000 bp (cb342 & cb343), 9.6 kb (cb1175 & cb1177 or cb343 & cb1338). Fluorescently labeled substrates were prepared by PCR amplification of pDR470 as follows: Cy3-100bp (cb1330 & cb1339), Cy3-9.6kb (cb1330 & cb1338), Cy5-100bp (cb1331 & cb1339), Cy5-9.6kb (cb1331 & cb1338). PCR products were purified by agarose gel electrophoresis.

Short double-stranded DNA substrates were prepared by annealing the following complementary oligonucleotides: 20 bp (cb939 & cb940), 40 bp (cb922 & cb935), 80 bp (cb95 & cb100). Substrates were labeled with [γ-^32^P]-ATP by polynucleotide kinase and gel purified. Binding reactions (20 µl) were carried out in 25 mM Tris-HCl pH 7.5, 7.5% glycerol, 100 mM NaCl, 2 mM DTT and 1 mg/ml BSA with 1 mM EDTA or 5 mM MgCl_2_, when indicated. Reactions contained 2 nM pUC19 or 0.5 nM radiolabeled substrate and the indicated concentration of protein. Concentrations for Rec114–Mei4 were calculated based on a 2:1 stoichiometry. For Mer2, the concentrations are expressed as monomers. Complexes were assembled for 30 minutes at 30 °C and separated by gel electrophoresis. For plasmid substrates, binding reactions were loaded on a 0.5% agarose (Gold) gel in 40 mM Tris-acetate buffer supplemented with 1 mM EDTA or 5 mM MgCl_2_, as indicated, at 50 V for 2.5 hours. Gels were stained with ethidium bromide and scanned using a ChemiDoc Imaging System (Bio-Rad). For short substrates, binding reactions were separated on 8% TAE-polyacrylamide gels at 200 V for 2 hours, gels were dried and imaged by autoradiography.

### In vitro condensation assays

DNA-driven condensation reactions were assembled as follows: RMM proteins were first diluted to 5 µl in storage buffer adjusted to a final salt concentration of 360 mM NaCl. After 5 minutes at room temperature, condensation was induced by 3-fold dilution in reaction buffer containing DNA and no salt, to reach final 15-µl reactions that contained 25 mM Tris-HCl pH 7.5, 5% glycerol, 120 mM NaCl, 2 mM DTT, 1 mg/ml BSA, 5 mM MgCl_2_, 5% PEG 8000, unless indicated otherwise. A typical binding reaction contained 150 ng pUC19 (5.7 nM), 50–200 nM Mer2 (^Alexa488^Mer2 or ^eGFP^Mer2) and/or 8–35 nM Rec114–Mei4 (^Alexa594^Rec114–Mei4 or ^mScarlet^Rec114–Mei4). After 30 minutes incubation at 30 °C with occasional mixing, 4 µl were dropped on a microscope slide and covered with a coverslip. Images were captured on a Zeiss Axio Observer Z1 Marianas Workstation with a 100×/1.4 NA oil immersion objective. Marianas Slidebook (Intelligent Imaging Innovations) software was used for acquisition. Images were analyzed with Image J using a custom-made script. Briefly, 129.24 × 129.24 µm (2048 × 2048 pixels) images were tresholded using the mean intensity of the background plus 3 times the standard deviation of the background. For experiments where the number of foci is compared between wild-type and mutant proteins or between reactions with and without Mg^2+^, a fixed threshold was applied. Masked foci were counted and the intensity inside the foci mask was integrated. Datapoints represent averages of at least 8–10 images per sample.

### Yeast strains and targeting vectors

Yeast strains were from the SK1 background. All strains used in this study are listed in **Supplementary Table S4**.

Strains that have endogenous *MER2* replaced by *kanMX4* cassette (SKY1524 and SKY1525) were described (Henderson et al., 2006). *MER2myc5::URA3* was inserted at the *mer2Δ::kanMX4* locus by EcoRI linearization of pRS306-derived pSK351 (WT) and pJX005 (KRRR) and transformation into SKY1524 and SKY1525 to yield SKY1560 and SKY1695 (WT), and SKY6411 and SKY6413 (KRRR). Integration of the vectors was confirmed by PCR.

Strains that have endogenous *REC114* replaced by the *kanMX4* cassette (SKY865 and SKY866) were described (Maleki et al., 2007). Tagged and untagged *REC114* alleles were generated by transformation of SKY865 and SKY866 with AflII-digested plasmids pRS305-derived targeting vectors. Plasmids and resultant strains were as follows: *REC114-8myc* (pSK591, SKY6749 & SKY6750), *REC114* (pSK592, SKY6562 & SKY6563), *rec114(F411A)-8myc* (pCCB857, SKY6889 & SKY6890), r*ec114(F411A)* (pCCB856, SKY6885 & SKY6886), *rec114(4KR)-8myc* (pCCB851, SKY6859 & SKY6860), *rec114(HLS)-8myc* (pSP113, SKY6797 & SKY6798).

Y2H vectors for wild-type DSB proteins were described previously (Arora et al., 2004; Maleki et al., 2007). pACT2-derived plasmids carry the *LEU2* marker and express the Gal4-activator domain. pCA1-derived plasmids carry the *TRP1* marker and express the DNA-binding domain of LexA. The vectors used here are as follows: pACT2-Rec114 (pSK304) encodes for Gal4AD-Rec114, pCA1-Mei4 (pSK281) encodes for LexA-Mei4, pCA1-Rec102 (pSK282) encodes for LexA-Rec102, pCA1-Rec104 (pSK283) encodes LexA-Rec104. Gal4AD empty vector control (pACT2) is pSK276. Y2H vectors for Rec114 truncations were generated by inverse PCR and self-ligation of the full-length construct pSK304. Plasmid numbers are as follows: Rec114(152-277) (pSP9), Rec114(del1-50&152-277) (pSP1), Rec114(del101-277) (pSP3), Rec114(del152-377) (pSP6). Rec114(53-428) and Rec114(1-377) were reported (Maleki et al., 2007). Point mutants were made by QuikChange mutagenesis and were as follows: Rec114-HLS (pSP25), Rec114-F411A (pCCB858).

### Immunofluorescence of yeast nuclei spreads

Diploid strains were cultured overnight in YPD (1% yeast extract, 2% peptone, 2% dextrose), followed by 13.5–14 hours in YPA (1% yeast extract, 2% peptone, 2% potassium acetate (KOAc)). Meiosis was induced by transfer to 2% KOAc. After 3.5 hours, cells were harvested, washed with H_2_O, resuspended in 1 M sorbitol, 1x PBS pH 7, 10 mM DTT, 0.5 mg/ml zymolyase 20T, and incubated for 30 minutes at 30 °C with gentle shaking. Spheroplasts were collected by centrifugation at 1500 g, washed in ice-cold 100 mM MES, 1 M sorbitol, spun down, then lysed in ice-cold 20 mM MES, 3 % paraformaldehyde and spread on a microscope slide for 1 hour. Slides were washed three times with 1 ml 0.4% PhotoFlo 200 solution (Kodak), air dried and stored at −20 °C. Slides were blocked with 90% FBS, 1× PBS for 1 hour at room temperature in a humid chamber, then incubated with primary antibody diluted in 3% BSA, 1× PBS in a humid chamber at 4 °C. After 3× 5-minute washes with 1× PBS in a Coplin jar, slides were incubated with secondary antibody diluted in 3% BSA, 1× PBS in a humid chamber at 37 °C for 1 hour. Slides were washed in the dark 3× 5 minutes with 1× PBS, mounted with Vectashield containing DAPI (Vector Labs). Primary antibodies used were mouse monoclonal anti-myc antibody clone 9E10 (1/100, Abcam) and rabbit polyclonal anti-Zip1 (1/50, this laboratory). Secondary antibodies used were goat anti-mouse IgG Alexa-488 and donkey anti-rabbit IgG Alexa-594 (1/200, Molecular Probes). Images of nuclei spreads were acquired on a Zeiss Axio Observer Z1 Marianas Workstation, equipped with an ORCA-Flash 4.0 camera and DAPI, FITC and Texas red filter sets, illuminated by an X-Cite 120 PC-Q light source, with a 100×/1.4 NA oil immersion objective. Marianas Slidebook (Intelligent Imaging Innovations) software was used for acquisition. Images were analyzed in Image J. Staging of nuclei spreads was based on DAPI staining and Zip1 immunufluorescence patterns, with nuclei showing a diffuse DAPI signal with either a single bright Zip1 focus or a few small Zip1 foci counted as leptotene or zygotene cells, respectively.

### Southern blot analysis of DSBs

Meiotic DSB analysis by Southern blotting was performed as described (Murakami et al., 2009). Briefly, synchronized cultures undergoing meiosis were harvested at the indicated time. After DNA purification, 800 ng of genomic DNA was digested by PstI and separated on a 1% TBE-agarose gel. DNA was transferred to Hybond-XL nylon membranes by vacuum transfer, hybridized with *SLY1* probe (amplified with primers: 5′-GCGTCCCGCAAGGACATTAG, 5′-TTGTGGCTAATGGTTTTGCGGTG) and developed by autoradiography.

### Spo11-oligo labeling

Procedure for labeling Spo11-associated oligonucleotides has been described (Neale and Keeney, 2009). Briefly, yeast cultures were harvested 4 hours into meiosis and denatured extracts were prepared by trichloroacetic acid precipitation. Proteins were solubilized in 2 % SDS, 500 mM Tris-HCl pH 8.1, 10 mM EDTA. Extracts were diluted in an equal volume of 2× IP Buffer (2 % Triton X100, 30 mM Tris-HCl pH 8.1, 300 mM NaCl, 2 mM EDTA, 0.02 % SDS) and Flag-tagged Spo11-oligo complexes were immunoprecipitated on IgG-conjugated agarose beads with mouse monoclonal M2 anti-Flag antibody. DNA was labeled on the beads with terminal deoxynucleotidyl transferase and [α-^32^P]-dCTP. After washing the beads in 1× IP buffer, proteins were eluted with LDS sample buffer and separated by SDS-PAGE. The gel was dried and developed by autoradiography.

### Western blotting of yeast meiotic extracts

Denaturing whole-cell extracts were prepared in 10% trichloroacetic acid with agitation in the presence of glass beads. Precipitated proteins were solubilized in Laemmli sample buffer and appropriate amounts of protein were separated by SDS-PAGE and analyzed by western blotting. Antibodies for western blotting were mouse monoclonal anti-myc (1/2000, Abcam), rabbit polyclonal anti-Kar2 (y-115) (1/2000, Santa Cruz). Secondary antibodies were used at 1/5000: IRDye 800CW goat anti-mouse IgG, IRDye 680 goat anti-rabbit IgG. Western blots were revealed using the Li-COR Bioscience Odyssey infrared imaging system.

### Yeast two hybrid

Y2H vectors were transformed separately in haploid strains SKY661 and SKY662 and selected on appropriate synthetic dropout medium. Strains were mated and streaked for single diploid colonies on medium lacking tryptophan and leucine. Single colonies were grown overnight in selective medium containing 2% glucose. Cultures were diluted in fresh medium containing 2% galactose and 1% raffinose and grown until log phase (4 hours). Cells were lysed and quantitative β-galactosidase assay was performed using ONPG substrate following standard protocols (Clontech Laboratories).

## End Matter

### Author Contributions and Notes

C.C.B. and S.K. designed research, C.C.B., S.P. and J.W. performed research, C.C.B, S.P., J.W. D.P. and S.K. analyzed data; and C.C.B. and S.K. wrote the paper.

The authors declare no conflict of interest.

## Acknowledgments

We thank Alain Nicolas for sharing unpublished information, Jiaqi Xu for assistance with preliminary analyses of the Mer2-KRRR mutant, and other members of the Keeney lab for discussions. We thank MSKCC core facilities, supported by NIH cancer center core grant P30 CA008748: Microchemistry and Proteomics (Ronald Hendrickson and Elizabeth Chang) for the XL-MS experiments, Molecular Cytology (Matthew Brendel and Yevgeniy Romin) for AFM experiments, and Sho Fujisawa for writing a Fiji script to quantify fluorescent foci. We thank Ewa Folta-Stogniew from the Biophysics Resource of Keck Facility at Yale University for the SEC-MALS experiments. The SEC-LS/UV/RI instrumentation was supported by NIH Award Number 1S10RR023748-01. The content of this paper is solely the responsibility of the authors and does not necessarily represent the official views of the National Institutes of Health. This work was funded by the Howard Hughes Medical Institute (SK) and an MSKCC Basic Research Innovation Award (SK and DP).

## Supplementary files

### Supplementary Figures

**Supplementary Figure 1:**
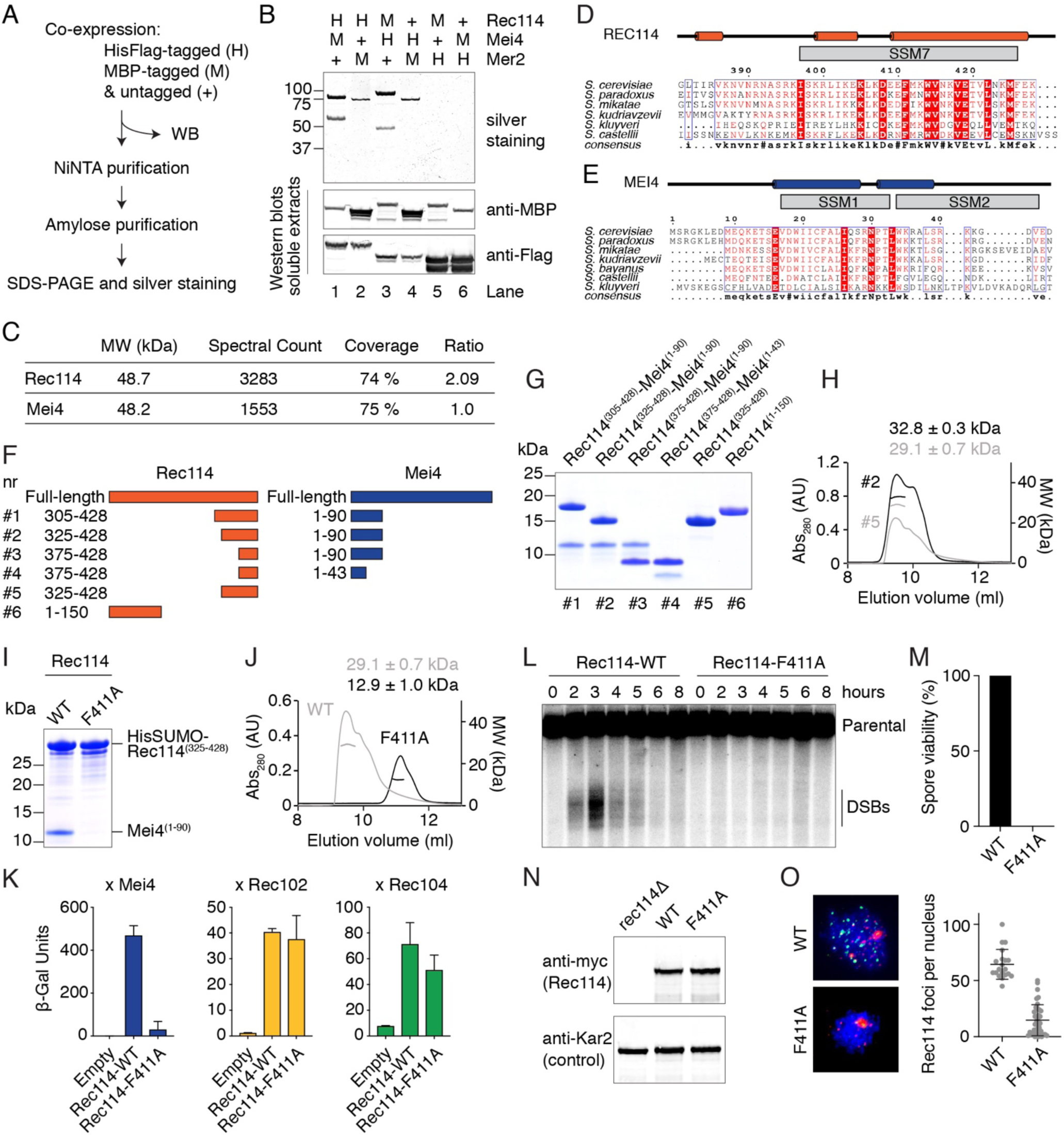
Characterization of the Rec114–Mei4 complex. A. Strategy for purification of a hypothetical Rec114–Mei4–Mer2 (RMM) complex. Combinations of MBP-tagged and HisFlag-tagged RMM subunits were co-expressed in insect cells. After cell lysis, complexes were purified by sequential affinity chromatography and analyzed by SDS-PAGE. Expression and solubility of the recombinant proteins are verified by western blotting (WB) of cell extracts. B. Analysis of purified complexes. Rec114– Mei4 complexes were apparent (lanes 1 and 3), but no Mer2 was co-purified. Lanes 2 and 4 show some enrichment of MBP-Mer2, but no co-purification of Rec114–Mei4. The presence of MBP-Mer2 in lanes 2 and 4 of the silver-stained gel may be due to background binding of MBP-Mer2 to the NiNTA resin (potentially via adsorption of DNA to the resin), or to low-affinity interactions to immobilized His-tagged Rec114–Mei4 complexes. Either way, none of the combinations tested yielded stoichiometric complexes of all three RMM subunits. Western blot controls of cell extracts showed that the tagged RMM proteins were expressed and soluble. C. Mass spectrometry analysis of Rec114–Mei4 complexes. Purified Rec114-Mei4 complexes were treated with trypsin and analyzed by LC-MS/MS. The ratio of spectral counts between Rec114 and Mei4 provides additional evidence supporting the 2:1 stoichiometry of the complex. D, E. Alignments and predicted secondary structures of the C-terminus of Rec114 (D) and the N-terminus of Mei4 (E). The positions of the conserved SSMs are indicated. F. Cartoon of the Rec114–Mei4 truncations analyzed. G. Purification of Rec114–Mei4 truncations. Proteins were expressed in *E. coli* and purified on NiNTA resin using a HisSUMO tag fused to the N-terminus of the Rec114 fragment. After removal of the tag by treatment with the SUMO protease Ulp1, complexes were further purified by gel filtration. A Coomassie-stained SDS-PAGE analysis of purified complexes is shown. 5 µg was loaded for each sample. Polypeptides containing Rec114^(375-428)^ and Mei4^(1-43)^ retained the ability to interact (combination #4). H. SEC-MALS analysis of Rec114–Mei4 truncations. The data are consistent with expectation for truncations that contain two Rec114 subunits and one Mei4 subunit. The C-terminus of Rec114 alone forms a dimer. I. Wild type and F411A-containing variants of ^HisSUMO^Rec114^(325-428)^ were co-expressed with Mei4^(1-90)^ and purified by chromatography on NiNTA resin. The absence of the Mei4 fragment with Rec114-F411A shows that the mutation abolishes the interaction with Mei4. J. SEC-MALS analysis of untagged wild-type (WT, reproduced from panel H to aid comparison) and F411A Rec114^(325-428)^ show that the mutation affects Rec114 dimerization. K. Y2H analysis of the interaction of Gal4BD-Rec114 (WT and F411A) with LexA-Mei4, LexA-Rec102, or LexA-Rec104. β-Gal units are quantified based on hydrolysis of ONPG. The F411A mutation abolishes the interaction of Rec114 with Mei4, but not with Rec102 and Rec104. Error bars represent SD from four replicates. L. Southern blot analysis of meiotic DSB formation at the *CCT6* hotspot, showing that *rec114-F411A* is defective in meiotic DSB formation. M. Spore viability of *rec114-F411A* mutant. N. Western-blot analyses of meiotic protein extracts from myc-tagged *REC114-WT* and *F411A* strains. The F411A mutation does not compromise the expression of Rec114. O. Immunofluorescence microscopy analysis of meiotic chromosome spreads with wild-type and F411A myc-tagged Rec114. Green, anti-myc; red, synaptonemal complex component Zip1; blue, DNA. Quantification of the number of Rec114 foci per leptotene or early zygotene cell is plotted; error bars show mean ± SD. The F411A mutation abolishes the formation of chromatin-associated Rec114 foci.

**Supplementary Figure 2:**
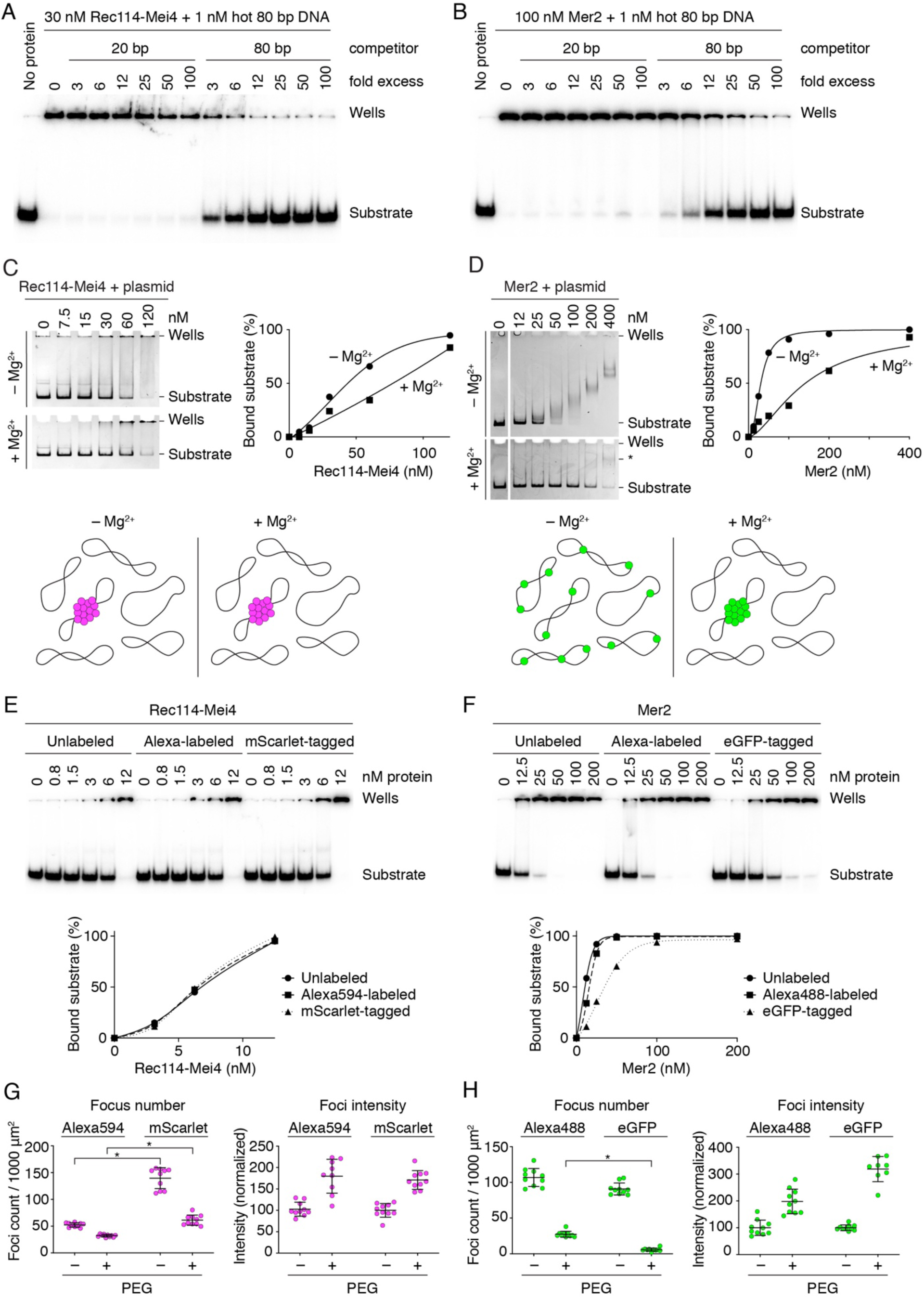
DNA-binding properties of Rec114–Mei4 and Mer2 complexes. A, B. Rec114–Mei4 and Mer2 bind with greater affinity to 80 bp substrates than 20 bp substrates. Competition assay of Rec114–Mei4 (A) and Mer2 (B) binding to an 80 bp radiolabeled DNA substrate (1 nM) in the presence of 20 bp or 80 bp cold competitor substrates. Fold excess is expressed in nucleotides. Quantification of these experiments is provided in **Figure 2D and F**. C, D. Binding of Rec114–Mei4 and Mer2 to plasmid DNA substrates analyzed by native agarose gel electrophoresis. Rec114–Mei4 (C) and Mer2 (D) were titrated with 2 nM plasmid DNA (pUC19) in the presence or absence of 5 mM MgCl_2_. Rec114–Mei4 complexes bound to plasmid DNA with roughly similar affinity independently of the presence of Mg^2+^ (apparent K_D_ ≈ 50–80 nM). Note that the apparent affinity is significantly lower than suggested by the gel shift analyses with radiolabeled substrates presented in **Figure 2A** (see apparent affinities in figure legend). We interpret that this difference is because the proteins coalesce on a small fraction of the plasmid molecules, as illustrated in the cartoon below. Indeed, bound plasmids remained trapped in the wells, which is consistent with cooperative assembly of large nucleoprotein structures. Because each plasmid substrate provides many more binding sites than the short oligonucleotide substrates in **Figure 2A**, a higher concentration of protein is required to reach complete binding of all of the plasmid molecules. In contrast to Rec114–Mei4, Mer2 showed efficient plasmid binding in the absence of Mg^2+^ in this assay (K_D_ = 30 ± 2 nM) but binding appeared to be considerably inhibited in the presence of Mg^2+^ (K_D_ ≈ 150 nM), as indicated by the persistence of unbound substrate at high protein concentrations. However, while the electrophoretic mobility of Mer2-bound plasmids decreased steadily as the concentration of Mer2 increased in the absence of Mg^2+^, no such steady progression was observed when Mg^2+^ was included. Instead, a minority of bound substrates shifted to a low-mobility species (labeled * in panel D, bottom), indicating that they were occupied by multiple Mer2 complexes. We interpret that, rather than inhibiting DNA binding, Mg^2+^ promotes cooperativity, in agreement with the fluorescence microscopy analysis (**Figure 2J**). Note that the difference in migration distance of the plasmid between the +/- Mg^2+^ gels is due to the presence of Mg^2+^ in the electrophoresis buffer. E–H. Effect of fluorophore labeling or tagging on the DNA-binding and DNA-driven condensation activities of Rec114–Mei4 and Mer2 complexes. Labeling with Alexa594 or Alexa488 was achieved using amine-reactive fluorophores. Tagging was achieved by fusion of Rec114 with the monomeric fluorescent protein mScarlet or fusion of Mer2 with the weakly dimerizing fluorescent protein eGFP. The results described here indicate that the covalent Alexa labeling has little if any effect on DNA binding properties of these complexes, whereas fluorescent protein tagging caused subtle alterations in DNA binding and/or condensation. E. Gel-shift analysis of binding of unlabeled, Alexa594-labeled, or mScarlet-tagged Rec114–Mei4 complexes to an 80-bp radiolabeled DNA substrate. The three versions of the Rec114–Mei4 complex have the same intrinsic DNA-binding activity. F. Gel-shift analysis of binding of unlabeled, Alexa488-labeled, or eGFP-tagged Mer2 complexes to an 80-bp radiolabeled DNA substrate. The DNA-binding activity of the Alexa-labeled Mer2 complex is nearly identical to the untagged protein, but the eGFP-tagged complex has 3.5-fold reduced DNA-binding activity. G. A comparison between Alexa-labeled and mScarlet-tagged Rec114–Mei4 complexes for DNA-driven condensation. Focus numbers (left graphs) and total fluorescence intensity within foci normalized to the no-PEG samples (right graphs) are shown for the complexes in the presence or absence of 5% PEG. With and without PEG, mScarlet-tagged Rec114–Mei4 produced more foci than the Alexa-labeled version. Because intrinsic DNA binding was indistinguishable between the complexes (panel E), we infer that the mScarlet-tagged complexes had a reduced efficiency in the cooperative formation of large condensates compared to the Alexa-labeled version, producing more numerous foci. Asterisk indicates p < 0.0001 (two-tailed t test). H. A comparison between Alexa-labeled and eGFP-tagged Mer2 complexes fpr DNA-driven condensation. Quantification is presented as in panel G. The two labeled complexes show different numbers and intensities of foci in the presence of PEG. It is likely that the DNA-binding defect of the eGFP construct (panel F) leads to the formation of fewer, brighter condensates. It is possible that the weak dimerization activity of eGFP also contributes. Asterisk indicates p < 0.0001 (two-tailed t test).

**Supplementary Figure 3:**
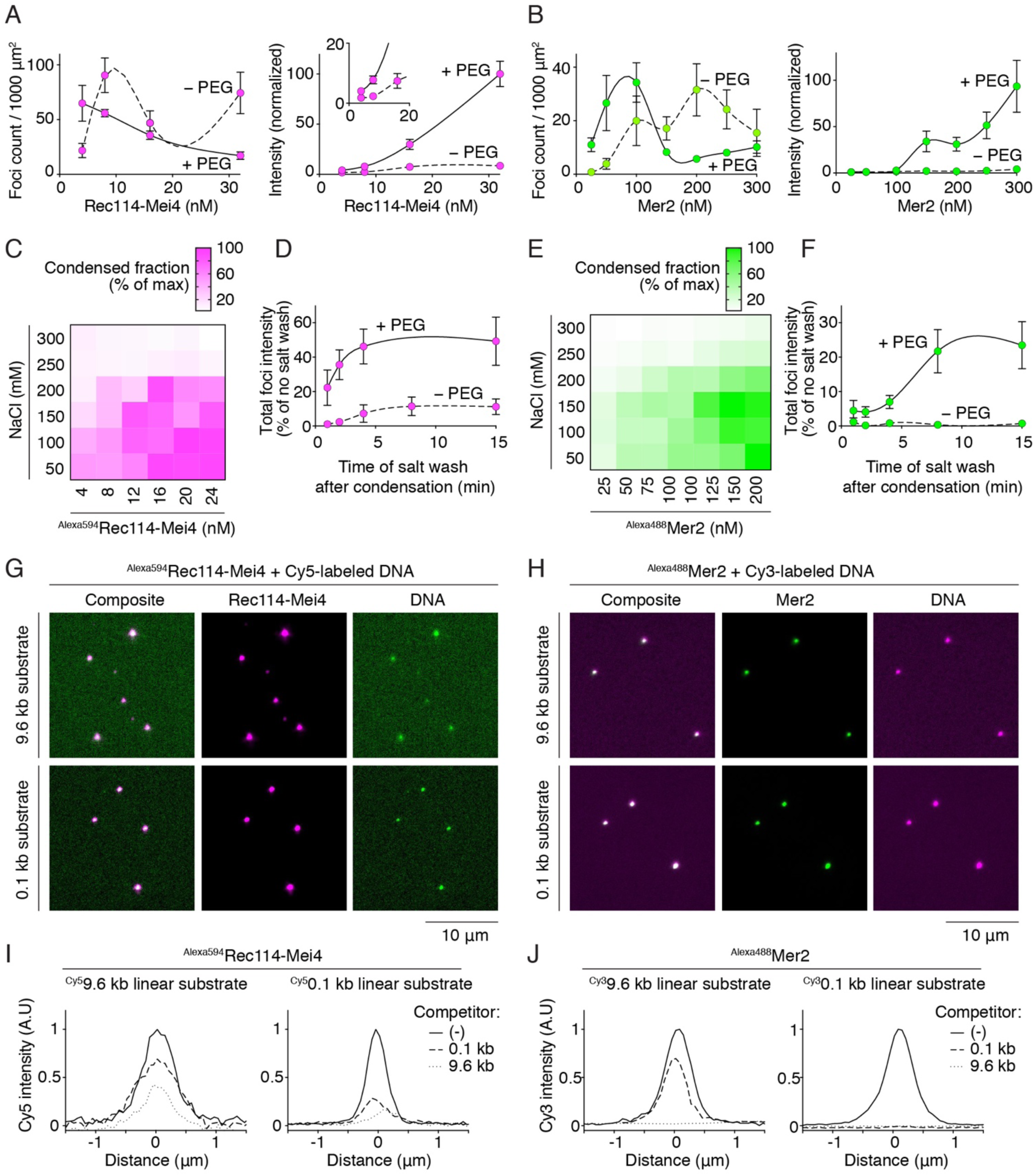
Properties of Rec114–Mei4 and Mer2 DNA-dependent condensates. A, B. Effect of Rec114–Mei4 (A) or Mer2 (B) concentration on DNA-driven condensation in the presence or absence of 5% PEG. Left graphs show focus numbers and right graphs show the total fluorescence intensity within foci (normalized to the mean of the highest intensity sample). Points and error bars are means ± SD from 8–10 fields of view. The titrations reveal complex behaviors: (A) In the presence of PEG, titration of Rec114–Mei4 from 4 to 32 nM led to a steady decrease in the number of foci, which was accompanied by a concomitant increase in focus intensity. In the absence of PEG, however, the number of Rec114–Mei4 foci first peaked at 8 nM before decreasing as the intensity of the foci started to increase. Nevertheless, focus intensity plateaued at a much lower intensity than in the presence of PEG. (B) In the case of Mer2, titration from 25 to 300 nM in the presence of PEG yielded a peak in the number of foci at ∼100 nM, which then sharply declined and stabilized beyond 150 nM. Consistently, Mer2 foci remained at a constant, low intensity between 25 and 100 nM, then became abruptly brighter above 100 nM. In the absence of PEG, the number of Mer2 foci increased between 25 and 200 nM, then started to decrease beyond that threshold. These behaviors likely reflect complex combined effects of nucleation, growth, and collapse of the condensates, which are each affected differently by protein concentrations and by the crowding effect provided by PEG. C, E. Titrations of Rec114–Mei4 (C) and Mer2 (E) in the presence of DNA and PEG and various concentrations of NaCl. Heat maps represent the fraction of fluorescence signal found within foci. Condensed fractions are maximal at high protein and low salt concentrations. At all protein concentrations, condensation is essentially abolished beyond 250 mM NaCl. This suggests that electrostatic interactions, likely between the negatively charged DNA backbone and positively charged protein residues, are important for condensation. D, F. Time dependence for irreversibility of Rec114– Mei4 (D) and Mer2 (F) condensates. Some phase-separated liquid droplets have been shown to mature over time and progressively adopt gel-like or solid states (Lin et al., 2015; Patel et al., 2015; Xiang et al., 2015; Banani et al., 2017). Such sol-gel transitions may occur spontaneously through different mechanisms, including fibrillization and entanglement, and are thought to be counteracted *in vivo* to prevent the progressive accumulation of amyloid-like structures associated with pathological states (Banani et al., 2017). To address whether our condensates are prone to progressive hardening, we queried the effect of assembly time on reversibility. We performed a time-course experiment where the condensates were challenged by treatment with 0.5 M NaCl after an indicated period of assembly in the presence or absence of PEG. The graph shows the total intensity summed for foci within fields of view, expressed as a percentage of the intensity without a salt challenge. Points and error bars are means ± SD for 8–10 fields of view. With Rec114–Mei4, 10% and 50% of fluorescent signal became refractory to the salt wash within 5 minutes of incubation time in the absence and presence of PEG, respectively (see **Figure 3C** for example images and quantification). With Mer2, there was no evidence for the formation of irreversible structures in the absence of PEG during the course of the experiment. However, up to 25% of the focus intensity resisted the salt wash treatment after 8 minutes of incubation time in the presence of PEG. Therefore, both Rec114–Mei4 and Mer2 have a propensity to form more stable, perhaps gel-like, structures over time. Under our experimental conditions, this was more evident for Rec114–Mei4 than for Mer2, and was accentuated by molecular crowding. G, H. Assembly of Rec114–Mei4 (G) and Mer2 (H) with fluorescently labeled 9.6 kb and 100 bp linear DNA substrates. The overlap between the protein foci and puncta of DNA shows that the DNA is also enriched in the condensates. However, in contrast to the protein signal, the fluorescent signal of the DNA covers the slide because DNA is in excess and does not condense by itself. I, J. Competition between long and short DNA substrates for incorporation into condensates. Rec114–Mei4 (I) or Mer2 (J) condensates were assembled in the presence of a fluorescently labeled DNA substrate with or without 20-fold nucleotide excess of unlabeled competitor. The amount of fluorescent DNA signal averaged between ten foci are plotted. In each case, the 9.6-kb substrate was a more effective competitor than the 100-bp substrate. In addition, the 100-bp substrate was more successful at competing with the 100-bp fluorescent substrate than with the 9.6-kb fluorescent substrate. This preference for large DNA substrates is consistent with the hypothesis that the condensates form through multivalent interactions between the positively charged residues of Rec114-Mei4 or Mer2 and the sugar-phosphate backbone of the DNA.

**Supplementary Figure 4:**
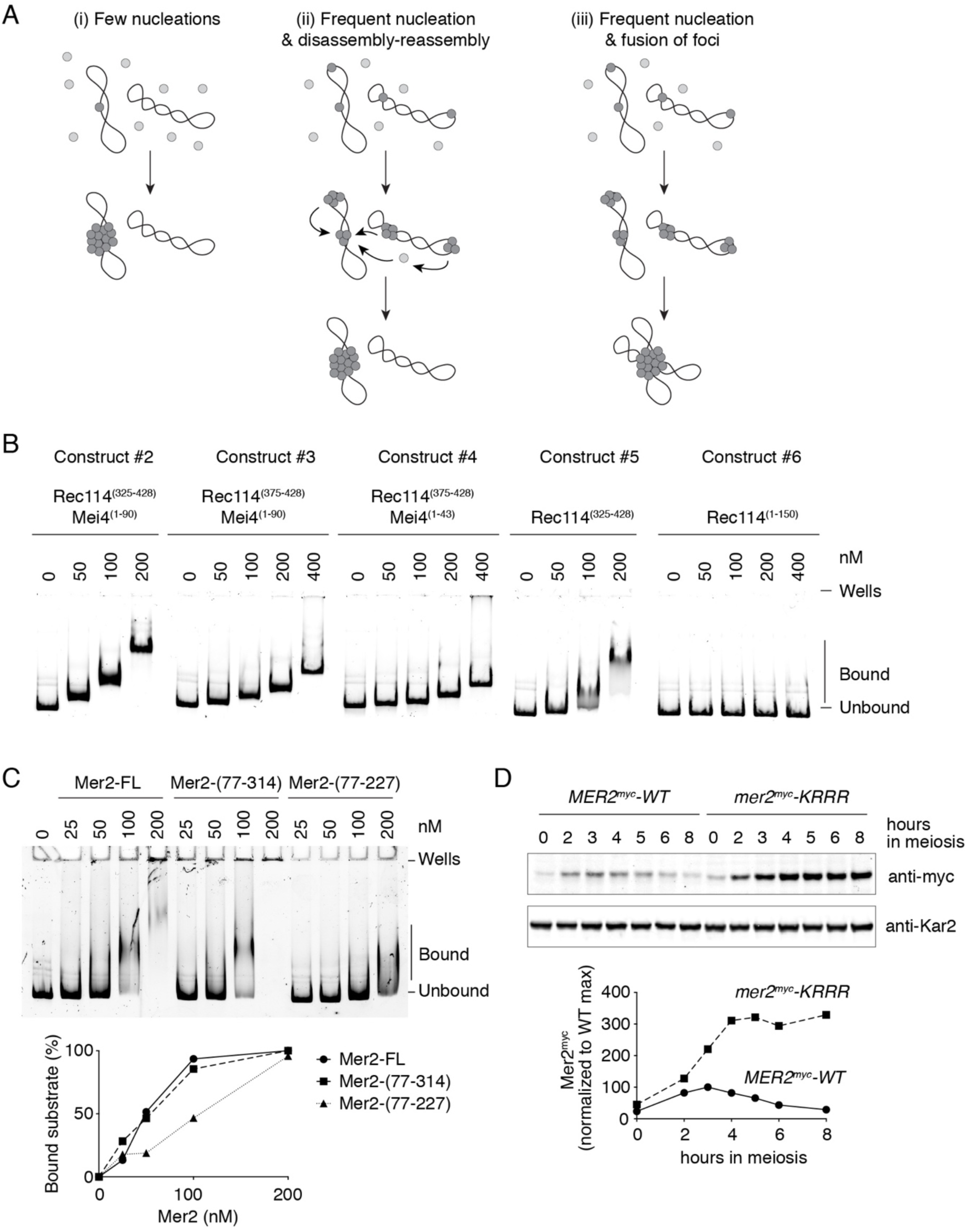
Mechanism of DNA-driven condensation and identification of DNA-binding residues. A. Three scenarios for the assembly of DNA-driven condensates (see text for details). B. Mapping the DNA-binding domain of Rec114–Mei4 complexes. Gel-shift analysis was performed with a pUC19 plasmid DNA substrate and the Rec114–Mei4 protein constructs shown in **Supplementary Figure 1F**. Constructs #2, #3 and #4, which include the C terminus of Rec114 and the N terminus of Mei4 were competent for DNA binding. The difference in mobility of shifted species between these constructs is in line with the difference in sizes of the protein complexes. Mei4 is dispensable for DNA binding by Rec114 (Construct #5 lacks Mei4). The N terminus of Rec114 alone, encompassing the PH domain, did not bind DNA (Construct #6). None of the constructs showed evidence for cooperative DNA binding (unlike the full-length protein, see **Supplementary Figure S2C**), suggesting that they do not undergo DNA-driven condensation. C. Mapping the DNA-binding domain of Mer2. Gel-shift analysis was performed with a pUC19 plasmid DNA substrate and HisSUMO-tagged Mer2 protein that was either full-length (FL), had the N terminus removed (fragment 77-314) or had both the N and C termini removed (fragment 77-227). Deleting the N terminus alone had no significant effect on DNA binding, but further deleting the C terminus strongly reduced DNA binding. D. Western blot analysis of Mer2-WT and Mer2-KRRR. Protein extracts of meiotic time courses were analyzed by SDS-PAGE followed by immunoblotting against Mer2-myc. Anti-Kar2 was used as a loading control. Quantification of western blot signal is plotted. Mer2^myc^-KRRR reached higher steady-state protein levels and persisted longer than wild-type Mer2^myc^. A previous study showed that mutating an essential CDK phosphorylation site of Mer2 (Ser30) or inhibiting CDK activity led to reduced turnover of Mer2, similar to the effect of the KRRR mutant (Henderson et al., 2006). This is consistent with the hypothesis that Mer2 turnover is tied to phosphorylation, which requires DNA binding.

**Supplementary Figure 5:**
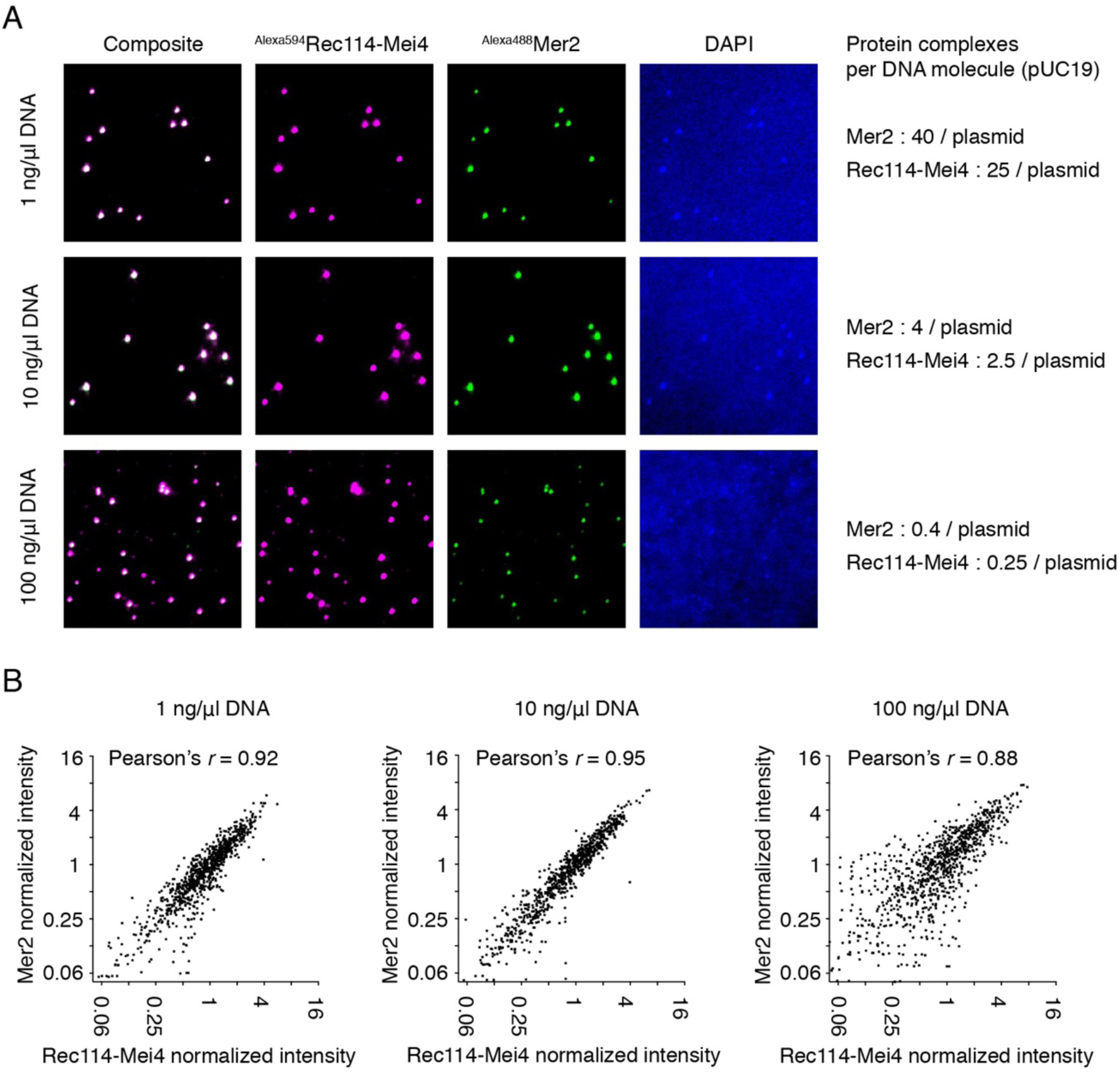
Rec114–Mei4 colocalizes with Mer2 in mixed condensates irrespective of DNA concentration. A. Reactions containing 16 nM Rec114–Mei4 and 100 nM Mer2 in the presence of 1, 10, or 100 ng/µl plasmid DNA were assembled for 20 minutes at 30 °C. DAPI (5 µg/ml) was added to the reaction before applying to glass slides. DNA enrichment within the condensates is visible at lower DNA concentrations (top and middle rows), but is not as clear at high DNA concentrations (bottom row). The ratios of Rec114–Mei4 (heterotrimers) and Mer2 (tetramers) to each 2.6-kb plasmid DNA molecule are indicated on the right. Colocalization of Rec114–Mei4 and Mer2 complexes is evident even with a molar excess of DNA molecules, demonstrating that formation of joint foci is not simply because both protein complexes are independently associating with a limiting number of DNA substrates. B. Correlated intensity of Rec114–Mei4 and Mer2 proteins within the condensates. Each point shows the fluorescence intensity in an individual focus (n > 900 foci from 2-3 fields of view), normalized to the average foci intensity per field of view. The strong correlation indicates that the composition of the condensates is highly uniform between foci. In the presence of high DNA concentration, the fraction of smaller foci increased and correlated intensities decreased.

### Supplementary Tables

Supplementary Tables 1: XL-MS data.

**Supplementary Table 2:**
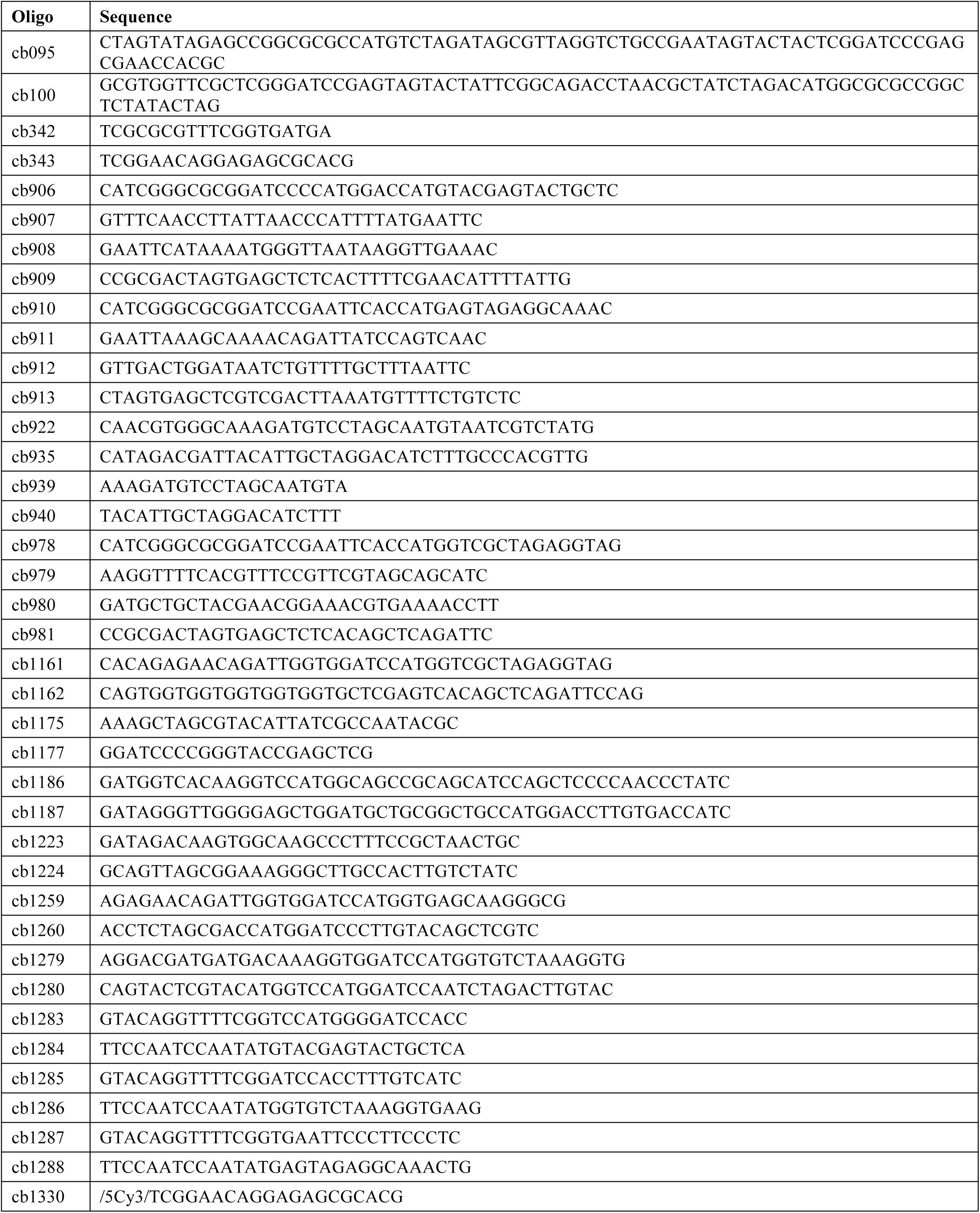
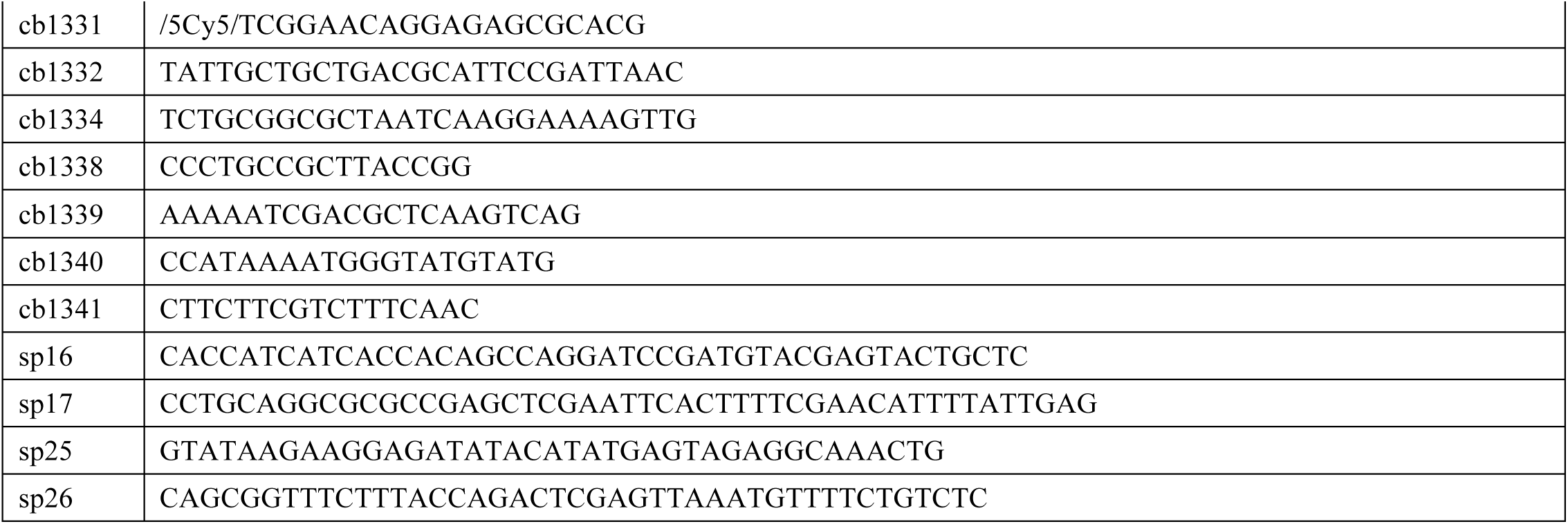
Oligonucleotides used in this study.

**Supplemental Table 3:** Plasmids used in this study.

| Plasmid | Description | Reference |
| --- | --- | --- |
| pCCB649 | Rec114 in pFastbac1 | This study |
| pCCB650 | HisFlag-Rec114 in pFastbac-HTb | This study |
| pCCB651 | MBP-Rec114 in pFastbac1 | This study |
| pCCB652 | Mei4 in pFastbac1 | This study |
| pCCB653 | HisFlag-Mei4 in pFastbac1-HTb | This study |
| pCCB654 | MBP-Mei4 in pFastbac1 | This study |
| pCCB681 | Mer2 in pFastbac1 | This study |
| pCCB682 | HisFlag-Mer2 in pFastbac1-HTb | This study |
| pCCB683 | MBP-Mer2 in pFastbac1 | This study |
| pCCB750 | HisSUMO-Mer2 in pSMT3 | This study |
| pCCB777 | HisSUMO-eGFP-Mer2 in pSMT3 | This study |
| pCCB779 | HisSUMO-Mer2-(K265A/R266A/R267A/R268A) in pSMT3 | This study |
| pCCB783 | HisSUMO-eGFP-Mer2-(K265A/R266A/R267A/R268A) in pSMT3 | This study |
| pCCB786 | HisFlag-mScarlet-Rec114 in pFastbac1 | This study |
| pCCB789 | HisFlag-Tev-Rec114 in pFastbac1 | This study |
| pCCB790 | HisFlag-Tev-mScarlet-Rec114 in pFastbac1 | This study |
| pCCB791 | MBP-Tev-Mei4 in pFastbac1 | This study |
| pCCB825 | HisSUMO-Rec114(375-428)-Mei4(1-43) in pETDuet1 | This study |
| pCCB835 | HisFlag-Tev-Rec114-(H39A/L40A/S41A) in pFastBac1 | This study |
| pCCB836 | HisFlag-Tev-mScarlet-Rec114-(H39A/L40A/S41A) in pFastBac1 | This study |
| pCCB848 | HisFlag-Tev-Rec114-(R395A/K396AK399A/R400A) in pFastBac1 | This study |
| pCCB849 | HisFlag-Tev-mScarlet-(Rec114-R395A/K396AK399A/R400A) in pFastBac1 | This study |
| pCCB850 | Rec114-(R395A/K396AK399A/R400A) in pRS305 | This study |
| pCCB851 | Rec114-(R395A/K396AK399A/R400A)-myc8 in pRS305 | This study |
| pCCB856 | Rec114-(F411A) in pRS305 | This study |
| pCCB857 | Rec114-(F411A)-8myc in pRS305 | This study |
| pCCB858 | Rec114-(F411A) in pSK304 | This study |
| pJX005 | Mer2myc5-K265A/R266A/R267A/R268A in pSK351 (pRS306) | This study |
| pSK276 | Gal4AD empty Y2H vector | Arora *et al.*, 2004 |
| pSK281 | LexA-Mei4 Y2H vector | Arora *et al.*, 2004 |
| pSK282 | LexA-Rec102 Y2H vector | Arora *et al.*, 2004 |
| pSK283 | LexA-rec104 Y2H vector | Arora *et al.*, 2004 |
| pSK304 | Gal4AD-Rec114 Y2H vector | Arora *et al.*, 2004 |
| pSK351 | Mer2-5myc in pRS306 | Henderson *et al.*, 2006 |
| pSK591 | Rec114-8myc in pRS305 | Maleki *et al.*, 2007 |
| pSK592 | Rec114 in pSR305 | Maleki *et al.*, 2007 |
| pSP1 | Rec114 del 1-50, del 152-277 in pSK304 | This study |
| pSP3 | Rec114 del 101-277 in pSK304 | This study |
| pSP6 | Rec114 del 152-377 in pSK304 | This study |
| pSP9 | Rec114 del 152-277 in pSK304 | This study |
| pSP25 | Rec114-(H39A/L40A/S41A) in pSK304 | This study |
| pSP34 | Rec114-Mei4 in pEtDuet1 | This study |
| pSP53 | HisSumo-Rec114-Mei4 in pEtDuet1 | This study |
| pSP113 | Rec114-(H39A/L40A/S41A)-8myc in pRS305 | This study |

**Supplementary Table 4:**
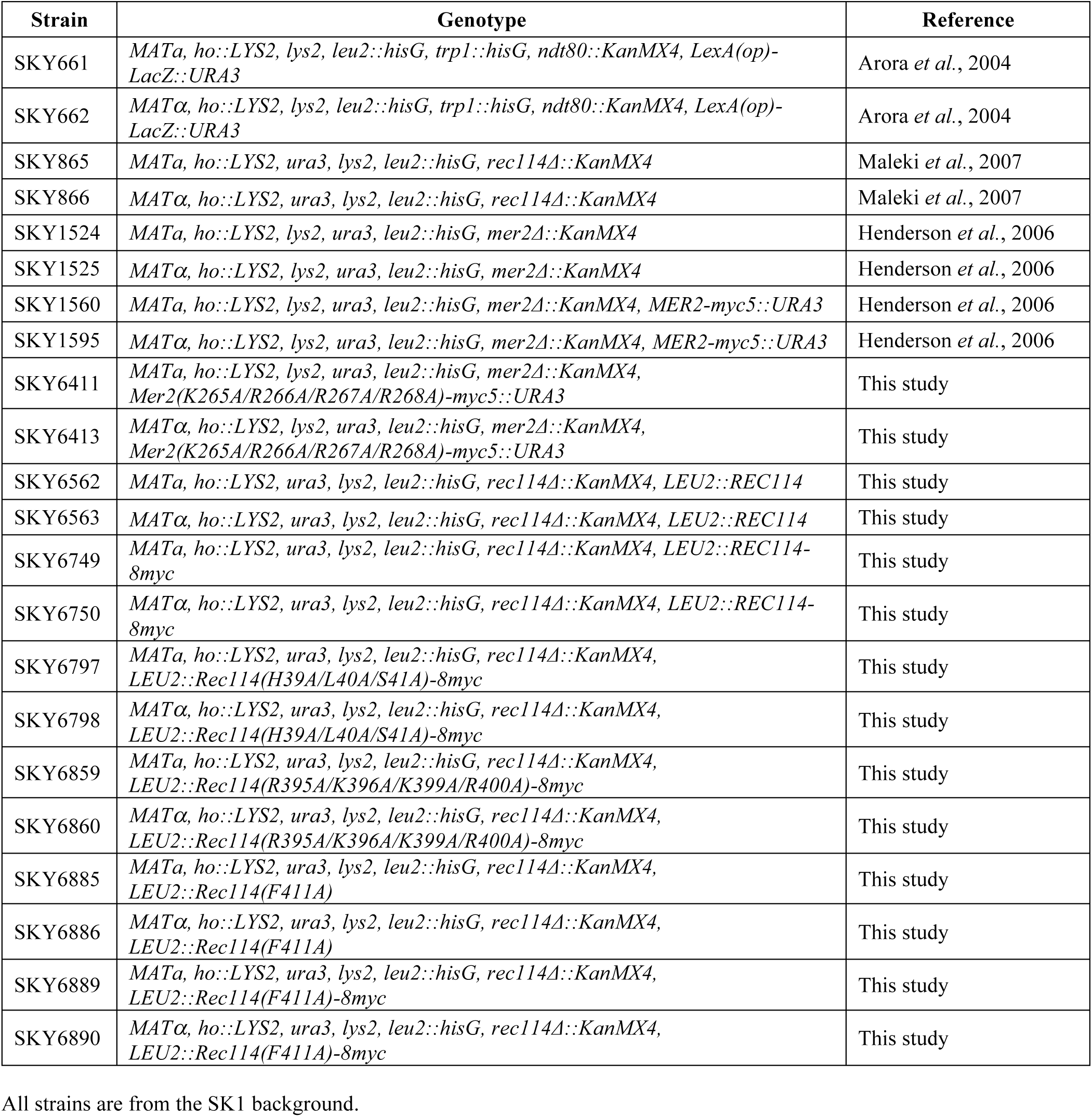
Yeast strains used in this study. All strains are from the SK1 background.

| Strain | Genotype | Reference |
| --- | --- | --- |
| SKY661 | *MATa, ho::LYS2, lys2, leu2::hisG, trp1::hisG, ndt80::KanMX4, LexA(op)-LacZ::URA3* | Arora *et al.*, 2004 |
| SKY662 | *MATα, ho::LYS2, lys2, leu2::hisG, trp1::hisG, ndt80::KanMX4, LexA(op)-LacZ::URA3* | Arora *et al.*, 2004 |
| SKY865 | *MATa, ho::LYS2, ura3, lys2, leu2::hisG, rec114Δ::KanMX4* | Maleki *et al.*, 2007 |
| SKY866 | *MATα, ho::LYS2, ura3, lys2, leu2::hisG, rec114Δ::KanMX4* | Maleki *et al.*, 2007 |
| SKY1524 | *MATa, ho::LYS2, lys2, ura3, leu2::hisG, mer2Δ::KanMX4* | Henderson *et al.*, 2006 |
| SKY1525 | *MATα, ho::LYS2, lys2, ura3, leu2::hisG, mer2Δ::KanMX4* | Henderson *et al.*, 2006 |
| SKY1560 | *MATa, ho::LYS2, lys2, ura3, leu2::hisG, mer2Δ::KanMX4, MER2-myc5::URA3* | Henderson *et al.*, 2006 |
| SKY1595 | *MATα, ho::LYS2, lys2, ura3, leu2::hisG, mer2Δ::KanMX4, MER2-myc5::URA3* | Henderson *et al.*, 2006 |
| SKY6411 | *MATa, ho::LYS2, lys2, ura3, leu2::hisG, mer2Δ::KanMX4, Mer2(K265A/R266A/R267A/R268A)-myc5::URA3* | This study |
| SKY6413 | *MATα, ho::LYS2, lys2, ura3, leu2::hisG, mer2Δ::KanMX4, Mer2(K265A/R266A/R267A/R268A)-myc5::URA3* | This study |
| SKY6562 | *MATa, ho::LYS2, ura3, lys2, leu2::hisG, rec114Δ::KanMX4, LEU2::REC114* | This study |
| SKY6563 | *MATα, ho::LYS2, ura3, lys2, leu2::hisG, rec114Δ::KanMX4, LEU2::REC114* | This study |
| SKY6749 | *MATa, ho::LYS2, ura3, lys2, leu2::hisG, rec114Δ::KanMX4, LEU2::REC114-8myc* | This study |
| SKY6750 | *MATα, ho::LYS2, ura3, lys2, leu2::hisG, rec114Δ::KanMX4, LEU2::REC114-8myc* | This study |
| SKY6797 | *MATa, ho::LYS2, ura3, lys2, leu2::hisG, rec114Δ::KanMX4, LEU2::Rec114(H39A/L40A/S41A)-8myc* | This study |
| SKY6798 | *MATα, ho::LYS2, ura3, lys2, leu2::hisG, rec114Δ::KanMX4, LEU2::Rec114(H39A/L40A/S41A)-8myc* | This study |
| SKY6859 | *MATa, ho::LYS2, ura3, lys2, leu2::hisG, rec114Δ::KanMX4, LEU2::Rec114(R395A/K396A/K399A/R400A)-8myc* | This study |
| SKY6860 | *MATα, ho::LYS2, ura3, lys2, leu2::hisG, rec114Δ::KanMX4, LEU2::Rec114(R395A/K396A/K399A/R400A)-8myc* | This study |
| SKY6885 | *MATa, ho::LYS2, ura3, lys2, leu2::hisG, rec114Δ::KanMX4, LEU2::Rec114(F411A)* | This study |
| SKY6886 | *MATα, ho::LYS2, ura3, lys2, leu2::hisG, rec114Δ::KanMX4, LEU2::Rec114(F411A)* | This study |
| SKY6889 | *MATa, ho::LYS2, ura3, lys2, leu2::hisG, rec114Δ::KanMX4, LEU2::Rec114(F411A)-8myc* | This study |
| SKY6890 | *MATα, ho::LYS2, ura3, lys2, leu2::hisG, rec114Δ::KanMX4, LEU2::Rec114(F411A)-8myc* | This study |

